# p53 controls choice between apoptotic and non-apoptotic death following DNA damage

**DOI:** 10.1101/2023.01.17.524444

**Authors:** Megan E. Honeywell, Marie S. Isidor, Nicholas W. Harper, Rachel E. Fontana, Peter Cruz- Gordillo, Sydney A. Porto, Cameron S. Fraser, Kristopher A. Sarosiek, David A. Guertin, Jessica B. Spinelli, Michael J. Lee

## Abstract

DNA damage can activate apoptotic and non-apoptotic forms of cell death; however, it remains unclear what features dictate which type of cell death is activated. We report that p53 controls the choice between apoptotic and non-apoptotic death following exposure to DNA damage. In contrast to the conventional model, which suggests that p53-deficient cells should be resistant to DNA damage-induced cell death, we find that p53-deficient cells die at high rates following DNA damage, but exclusively using non-apoptotic mechanisms. Our experimental data and computational modeling reveal that non-apoptotic death in p53-deficient cells has not been observed due to use of assays that are either insensitive to cell death, or that specifically score apoptotic cells. Using functional genetic screening – with an analysis that enables computational inference of the drug-induced death rate – we find in p53-deficient cells that DNA damage activates a mitochondrial respiration-dependent form of cell death, called MPT-driven necrosis. Cells deficient for p53 have high basal respiration, which primes MPT-driven necrosis. Finally, using metabolite profiling, we identified mitochondrial activity-dependent metabolic vulnerabilities that can be targeted to potentiate the lethality of DNA damage specifically in p53-deficient cells. Our findings reveal how the dual functions of p53 in regulating mitochondrial activity and the DNA damage response combine to facilitate the choice between apoptotic and non-apoptotic death.

## INTRODUCTION

Understanding the mechanisms of drug action is important for identifying settings in which a drug will be effective, and for interpreting, even predicting, potential mechanisms of drug resistance. For anti-cancer drugs, a key aspect towards this end is to understand the mechanism(s) by which these drugs promote cell death. The most well-studied form of cell death is apoptosis, and many anti-cancer drugs are known to function by activating apoptotic death^1,2^. In recent years, however, more than a dozen non-apoptotic forms of regulated cell death have been identified^3–6^. While these mechanisms may not be used in normal development, many non-apoptotic death pathways are valuable therapeutic targets due to their hyperactivation in pathological states, such as in some cancer cells^7^. Furthermore, some anticancer drugs can activate both apoptotic and non-apoptotic cell death, but how the choice is made in a particular context remains unclear^8,9^.

The DNA damage response (DDR) is a kinase-driven signaling pathway underlying the efficacy of most conventional chemotherapeutics^10,11^. In addition to recruiting DNA repair machinery, the DDR coordinates the selection of possible downstream cell fates, including cell cycle arrest, permanent senescence, or alternatively, activation of apoptotic cell death. In general, the functional outcome of DDR signaling is dictated by the dynamics of the tumor suppressor p53^12–15^. While the mechanisms by which p53 signaling and p53 dynamics dictate cell fate choices are becoming increasingly understood, it remains unclear how cell fate decisions following DNA damage are regulated in the absence of p53. Furthermore, because p53 is mutated or deleted in most cancers, understanding how p53-deficient cancer cells respond to DNA damage is critical for our understanding of anti-cancer drug action, and our ability to select companion therapies to improve responses.

Most prior studies have found that p53 is required for DNA damage-induced cell death^16–18^. These studies are well-validated and consistently reproducible; however, many DNA-damaging chemotherapies are effective in cancers that lack p53. For instance, medullary triplenegative breast cancers are universally mutant for p53, but paradoxically, these are among the most chemo-sensitive breast cancers^19,20^. Because cell death is generally thought to be required for durable disease remission^21^, and because most prior studies have focused exclusively on apoptotic death, we reasoned that other mechanisms must exist to facilitate DNA damage-induced cell death in the absence of p53. Indeed, DNA damage has been demonstrated to activate some forms of non-apoptotic death^22–25^, but the mechanisms of activation and the contexts that induce apoptotic versus non-apoptotic death remain unclear. Furthermore, because most forms of non-apoptotic death have only recently been recognized, this question remains largely unexplored.

We find that following DNA damage, p53 is not required for cell death, *per se*, but is specifically required for the activation of cell intrinsic apoptosis. Cells die at similar levels following DNA damage with and without p53, but p53-deficient cells preferentially activate a non-apoptotic form of cell death. Using functional genetic screening of single gene knockouts, complemented with a new analysis strategy to infer perturbations to the drug-induced death rate, we find that DNA damage activates a mitochondrial respiration-dependent form of cell death, called MPT-driven necrosis, in p53-deficient cells. Cells deficient for p53 are primed for MPT-driven necrosis due to hyperactive respiration and the formation of mitochondrial electron transport chain (ETC) supercomplexes (SC). Inhibiting respiration inhibits MPT-driven necrosis in p53-deficient cells, but does not alter DNA damage-induced death in p53-proficient cells. Additional metabolomic analyses reveals that the altered metabolism in p53-deficient cells creates a unique vulnerability to high NAD+ levels. Furthermore, nicotinamide mononucleotide (NMN) supplementation, which increases NAD+ production, sensitizes p53-deficient cells to DNA damage. Taken together, these findings reveal how the dual functions of p53 in regulating mitochondrial activity and the DDR combine to facilitate the choice between apoptotic and nonapoptotic death following DNA damage.

## RESULTS

### p53-deficient cells undergo non-apoptotic cell death following DNA damage

Much is known about how p53 contributes to the DDR; however, it is less clear how cell fate decisions following DNA damage are regulated in the absence of p53 (Figure 1A)^10^. To study this issue further, we began by profiling sensitivity to nine commonly used DNA-damaging chemotherapeutics. Each drug was tested in cells containing wild-type p53 (WT) or cells that lack p53 function (KO). We used the FLICK assay (**<u>F</u>**luorescence-based and **<u>L</u>**ysis-dependent **<u>I</u>**nference of **<u>C</u>**ell death **<u>K</u>**inetics), to quantify the numbers of live and dead cells, and how these populations change over time following drug addition^9,26^. To analyze these data, we focused on the drug-induced lethal fraction (LF; *i.e.*, percentage of the population that is dead, following drug exposure), or its inverse, fractional viability (FV). Unlike more conventional metrics, such as “relative viability” (RV; *i.e.*, relative number of live cells, compared to untreated control), LF and FV are cell death-specific measurements, whereas RV conflates growth inhibitory and death activating phenotypes^27^. Overall, we found no statistically significant differences in DNA damage-induced lethality between p53 WT and p53 KO cell lines (Figure 1B and S1A-B). These results are somewhat unexpected, considering the common interpretation that p53 is required for robust activation of apoptosis following DNA damage^17,18,28^. Nonetheless, similar results were also found following re-analysis of the DepMap PRISM Repurposing dataset, which contains 90 compounds annotated as DNA damaging agents, each profiled in approximately 400 cell lines (Figure 1C).

**Figure 1:**
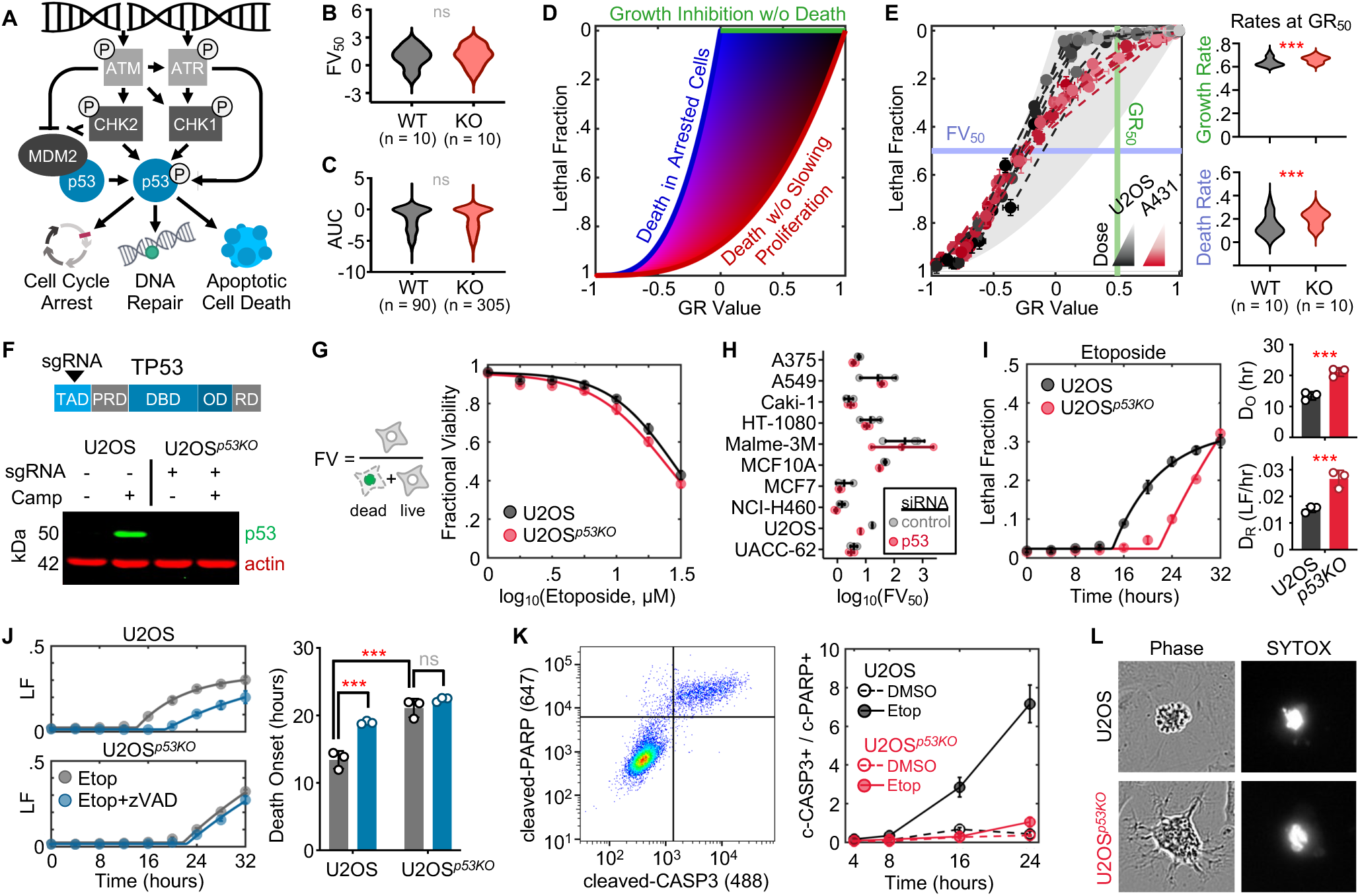
p53 deletion switches the mechanism of cell death following DNA damage from apoptotic to non-apoptotic. **(A)** Simplified schematic of the DNA damage response. **(B – C)** DNA damage sensitivity for p53-proficient (WT) and p53-deficient (KO) cell lines. (B) Sensitivity to 9 DNA damaging chemotherapeutics in p53 WT or KO cells. Data were generated using the FLICK assay, and scored using the IC_50_ of the fractional viability dose response (FV_50_). (C) Chemosensitivity as in (B), from the DepMap drug repurposing dataset. **(D)** Schematic of drug GRADE analysis. **(E)** Example GRADE analysis for U2OS (p53 WT) and A431 (p53 KO) treated with 9 DNA damaging drugs. Growth rates and death rates at GR_50_ effect size shown for the full panel of 10 WT and 10 KO cells in panel B. **(F)** Generation of U2OS*^p53KO^* cells. **(G)** Etoposide sensitivity of U2OS and U2OS*^p53KO^.* FV measured using FLICK. **(H)** FV_50_ for 10 p53 WT cells treated with etoposide in the presence and absence of p53-targeted siRNA. **(I)** Cell death kinetics for U2OS and U2OS*^p53KO^* treated with 31.6 µM etoposide. **(J)** As in panel I, but ± zVAD. **(K)** Apoptotic death evaluated using flow cytometry. Example for U2OS treated with etoposide (*left*), quantified (*right*). **(L)** Death morphology in U2OS and U2OS*^p53KO^.* Apoptotic “blebbing” morphology shown for U2OS. Non-apoptotic morphology in U2OS*^p53KO^.* SYTOX positivity reports loss of membrane integrity. For all panels with error bars, data are mean ± SD from 3 experimental replicates. See also Figure S1 and S2.

A potential issue with interpreting the role of p53 from a meta-analysis performed across drugs and cell lines is that p53 is thought to have functions that vary depending on context^29,30^. Thus, any critical role for p53 in DNA damage-induced cell death may be diluted by integrating across fundamentally different cellular and drug contexts. Additionally, dose summary metrics, such as the IC50 or FV50, may not capture variation in drug sensitivity that exists at other doses. To address these issues, and to more accurately score how p53 affects cell death following DNA damage, we used the drug GRADE framework to evaluate responses to DNA-damaging chemotherapies^27^. The GRADE approach derives from the comparison of two complementary measurements: the normalized net population growth rate (GR value^31^) and the drug-induced lethal fraction (Figure 1D)^32,33^. These measurements are normalized for growth rate variations between cell types, enabling a more accurate comparison of responses across different cellular contexts. Importantly, GRADE reveals the degree of coordination between growth inhibition and cell death activation, and how these features vary across doses of a drug.

Within any given cell line, GRADEs for DNA-damaging chemotherapies were generally similar, even across mechanistically distinct classes of DNA damage (Figure S1C). Importantly, GRADE-based analysis revealed striking differences between p53-proficient and p53-deficient cells in the coordination of growth inhibition and death activation. At high lethal doses (such as the FV50, shown in Figure 1B), all cell types have similar coordination, with death occurring only following complete growth arrest within the population. However, at lower doses, such as the GR50 (*i.e.,* dose in which the net population growth rate is reduced by half), distinct responses were observed between p53-proficient and –deficient cells. For instance, in U2OS, which have functional p53, the observed response at the GR50 dose is entirely the result of growth inhibition, with no activation of cell death (Figure 1E). In contrast, in A431, which are p53-deficient, cells continue to proliferate at the GR50 dose, but also activate cell death at high levels (Figure 1E). Similar results were found in a larger panel of p53 WT and p53 KO cells (Figure 1E and S1C). Thus, these data show that DNA damage activates cell death at *higher –* not lower – rates in p53-deficient cells, but these high death rates are compensated for by significantly higher growth rates. Similar results were observed at all effect sizes, and we failed to identify any dose in which DNA damage was more lethal in p53-proficient cells than in p53-deficient cells. Continued proliferation of p53-deficient cells following DNA damage is perhaps expected, given the role for p53 in facilitating DNA damage-induced G1-S cell cycle arrest^34^. However, given numerous studies which demonstrate that p53 is required for DNA damage-induced cell death, it is surprising that cells lacking functional p53 activate cell death at significantly higher rates than cells with functional p53. Thus, GRADE-based analysis reveals hidden and unexpected variation in the DNA damage responses of p53-proficient and p53-deficient cell lines. Furthermore, in contrast to the prevailing model, our data reveal that cells lacking p53 die at high rates following DNA damage.

The insights generated by our GRADE-based evaluation of drug responses are in apparent conflict with the well-validated model that p53 is required for apoptotic cell death following DNA damage. We reasoned that there were at least two possible parsimonious explanations for these unexpected results. First, tumor cells that evolve without functional p53 have higher levels of genome instability and a lower capacity for DNA repair^35^. Thus, these cells may be experiencing higher levels of DNA damage per dose, compared to cells with functional p53. A second possibility is that cells lacking p53 die primarily via non-apoptotic mechanisms. In the context of DNA damage, the preponderance of published evidence suggests that p53 is not required for the activation of cell death, *per se,* but specifically required for activation of apoptosis^28^. Furthermore, because cell death has typically been evaluated using markers that are specific to apoptotic cells (*e.g*., caspase-3 cleavage, Annexin V positivity, membrane blebbing), high levels of non-apoptotic death can occur and go unnoticed. To address these issues, we used CRISPR/Cas9-mediated genome editing to delete p53, facilitating an analysis of p53 function in an isogenic pair of cell lines (Figure 1F). We used U2OS cells, which have wild-type p53 and retain p53-dependent functions, such as p53-dependent G1-S arrest following DNA damage (Figure S2A-B). Loss of p53 did not substantially alter levels of DNA damage, or the duration of DNA damage signaling (Figure S2C). To evaluate the role of p53 in DNA damage-induced cell death, we tested the topoisomerase II inhibitor etoposide. Etoposide-induced death was similar for U2OS and U2OS*^p53KO^* cells, with the level of cell death modestly (but significantly) increased in the absence of p53 (Figure 1G). Similarly, acute knockdown of p53 using targeted siRNAs did not strongly alter the level of cell death in a panel of genetically diverse cells containing functional p53 (Figure 1H).

While the overall levels of cell death were similar in the presence and absence of p53, there were clear differences in the kinetics of drug-induced cell death between U2OS and U2OS*^p53KO^* cells. In particular, the onset time of cell death (D_O_) was significantly later in U2OS*^p53KO^*when compared to U2OS cells, but also death occurred at a faster overall rate (D_R_), once initiated (Figure 1I). Faster population death rates are an indication of increased synchrony of death within the population, which is a common feature of inflammatory non-apoptotic forms of death^33,36–38^. Thus, we also tested whether DNA damage activated apoptotic or non-apoptotic death in these cells. In response to etoposide, death in U2OS cells was associated with the conventional hallmarks of apoptotic death: sensitivity to the caspase inhibitor zVAD, activation of the apoptotic executioner, caspase-3, and acquisition of the characteristic “membrane blebbing” morphology (Figure 1J-L). These hallmarks were absent in U2OS*^p53KO^*cells, despite the high levels of death induced by DNA damage in these cells (Figure 1J-L).

An additional distinction between apoptotic and non-apoptotic death is that non-apoptotic forms of cell death often result in release of inflammatory mediators, whereas apoptosis is typically considered to be immunologically silent. Thus, we evaluated whether DNA damage-induced cell death in the presence and absence of p53 induced inflammatory responses in surviving cells. Conditioned media taken from U2OS*^p53KO^* cells treated with etoposide induced inflammatory responses, which were not observed from conditioned media taken from U2OS cells treated with etoposide (Figure S2D-F). A possible explanation for the lack of apoptosis in U2OS*^p53KO^* cells could be that these cells simply have a reduced ability to activate apoptosis. However, using the BH3 profiling assay we found that U2OS*^p53KO^* cells have modestly higher levels of apoptotic priming than U2OS (Figure S2G-H), and both cell types have similar sensitivities to the BH3 mimetic compound ABT-199 (Figure S2I). Thus, although DNA damage activates different types of cell death in U2OS and U2OS*^p53KO^* cells, these two cell types have similar capacities to activate apoptosis. Taken together, these data reveal that DNA damage activates cell death to similar levels in the presence and absence of p53, but via distinct mechanisms: cells with functional p53 die via apoptosis, whereas cells without functional p53 die via non-apoptotic mechanisms.

### Assay conditions and analysis methods prevented the identification of p53 independent cell death

Considering the intense interest in both p53 and cellular responses to DNA damage, we next sought to explore why it had not previously been recognized that DNA damaging agents cause high levels of non-apoptotic death in the absence of p53. Relative viability (RV) has overwhelmingly been the most common method for evaluating drug responses in all contexts, including p53-dependent responses to DNA damage. p53-deficient cells appear to have reduced sensitivity to DNA damage if evaluated using the common RV metric (Figure 2A). However, it has recently been clarified that evaluating relative drug sensitivities using RV-based analysis can be confounded by differences in proliferation rates between cell types^31,39^. Because p53 can regulate both proliferation and cell death, we reasoned that varied growth rates in the presence and/or absence of DNA damage likely contributed to the inability to accurately interpret the extent to which p53 regulates cell death following DNA damage.

**Figure 2:**
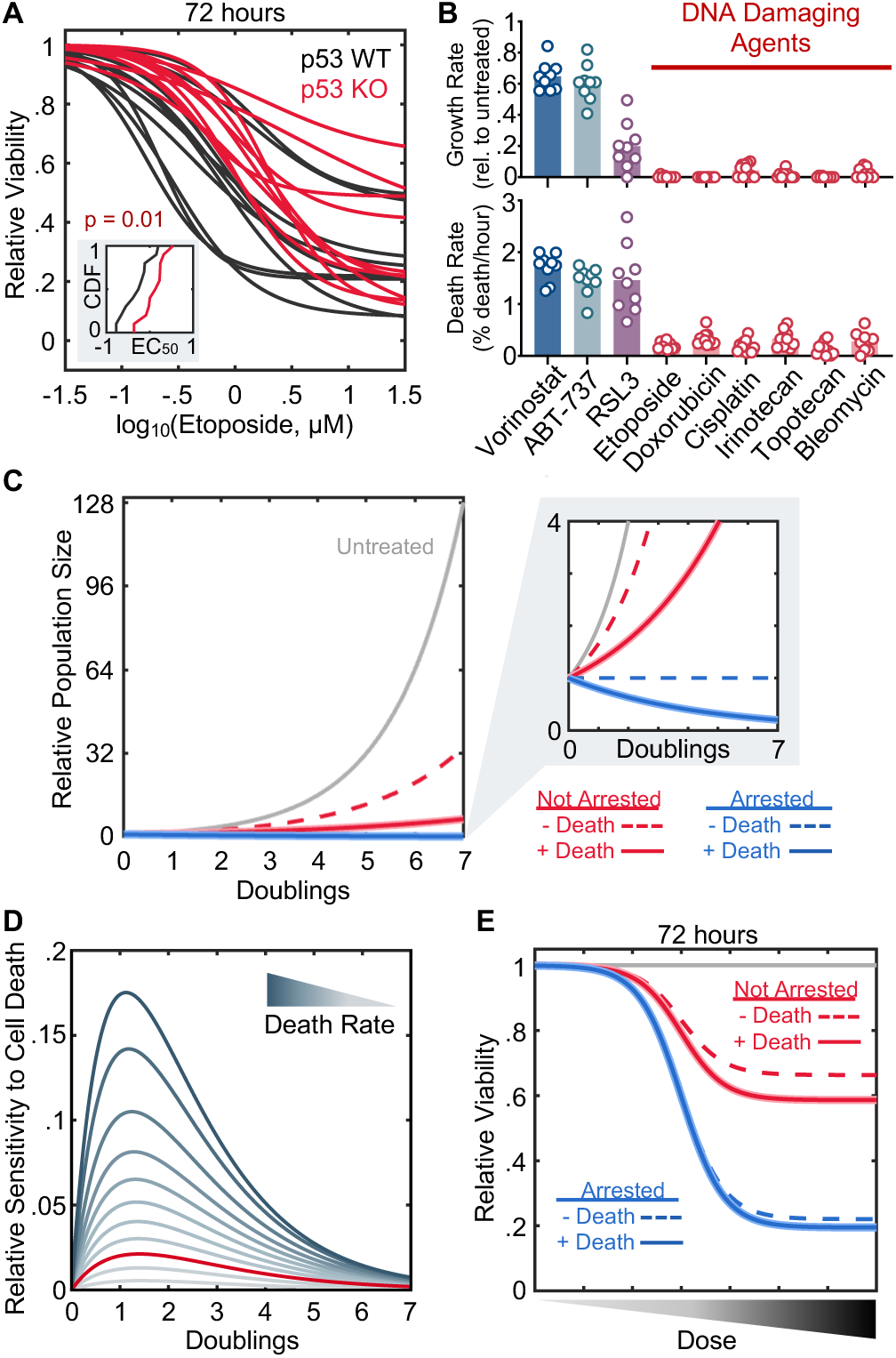
Use of alternative death mechanisms in the absence of p53 has been missed due to assay conditions that are insensitive to cell death. **(A)** Etoposide sensitivity for 10 p53 WT and 10 p53 KO cell lines, evaluated using Relative Viability. Inset shows cumulative distribution function (CDF) for EC_50_ in p53 WT and KO cells. p-value calculated using KS test. **(B)** Growth rates and death rates for HDAC inhibition (Vorinostat), BH3 mimetic (ABT-737), or a ferroptotic drug (RSL3), compared to 6 DNA damaging chemotherapeutics. Data were generated using the FLICK assay with GRADE analysis. **(C-E)** Simulation of population dynamics following exposure to DNA damage. **(C)** Relative number of live cells over time for different growth and death models (for cells with/without arrest, and with/without death following DNA damage). Growth and death rates are parameterized from observed rates in p53 WT and KO cells. **(D)** Sensitivity of the RV metric to cell death. The expected change in RV at different points in time, for simulations of drugs with different death rates. The peak sensitivity depends on the death rate, but RV universally becomes insensitive to the death rate over time. Red curve is the average death rate observed for DNA damaging agents. **(E)** Simulated RV dose-response function for cells with/without arrest, and with/without death following DNA damage. RV is much more sensitive to changes in proliferation rates than changes in death rates. See also Figure S2.

To consider this issue from a quantitative perspective, we built a simple model to simulate the population response to DNA damage. We used this model to quantitatively evaluate the expected changes in RV if p53 controls both the proliferation rate and death rate following DNA damage, or alternatively, if these phenotypes are uncoupled and p53 regulates the proliferation rate but not the death rate, as we see in our empiric experimental data. We parameterized our model using growth rates and death rates observed in our GRADE-based analysis (Figure 1 and S2B). In the presence of p53, DNA damage induces a bi-phasic response, characterized by growth inhibition at low doses, and cell death occurring only at higher doses and only in non-proliferative cells^27,40^. As a result, the experimentally observed death rate following high levels of DNA damage is surprisingly low, and this death rate would not be sufficient to shrink a tumor population, if the tumor cells were not also growth arrested (Figure 2B).

Our simulations revealed that loss of either p53-dependent growth inhibition or p53-dependent death activation should result in a similar *qualitative* phenotype of increased RV (dotted lines are always higher than solid lines, Figure 2C). Importantly, however, these phenotypes are only clearly observable at early times following drug exposure. Over time, the RV metric becomes insensitive to the drug-induced death rate as exponential proliferation overwhelms the RV scale (Figure 2D). At time points that are commonly used for drug evaluation studies, RV is ostensibly an exclusive evaluation of the drug induced-growth rate: in both a growing or arrested population, the presence or absence of drug-induced cell death does not strongly alter the overall population size (Figure 2E). Thus, due to a combination of three features – low intrinsic death rates for lethal doses of DNA damage, long assay times that bias towards anti-proliferative phenotypes, and common use of relative viability to measure drug sensitivity – the degree to which p53 does-or does not contribute to DNA damage-induced cell death would not be easily observed in prior studies.

### Regulators of DNA damage-induced non-apoptotic death identified using a death rate-focused analysis

Having found that cells lacking p53 die predominantly using non-apoptotic mechanisms following DNA damage, we next sought to determine which form of non-apoptotic death was being activated. At least 15 mechanistically distinct subtypes of cell death have been characterized, most of which induce a morphologically necrotic form of death^3–6,41^. We began by evaluating eight pathways, for which we had well-validated inhibitors. Single pathway inhibitors failed to inhibit the lethality of etoposide in U2OS*^p53KO^* cells, as did higher order combinations of these inhibitors (Figure S3A-B). Thus, in the absence of p53 it appears that death following DNA damage occurs using either a poorly characterized form of cell death, or a combination of death subtypes that we were unable to evaluate using chemical inhibitors.

We next sought to identify which subtype of non-apoptotic death is activated in p53-deficient cells using functional genetics. We used the GeCKOv2 sgRNA library to generate single gene knockouts, in Cas9-expressing U2OS or U2OS*^p53KO^* cells treated with the topoisomerase II inhibitor, etoposide (Figure 3A)^42^. In most prior studies, drugs have been screened at doses that confer a 20-50% reduction in population size over 2-4 weeks (*e.g.*, “ED20” dose; approximately 1 nM for etoposide)^43^. These doses of etoposide, however, fail to induce any cell death in U2OS or U2OS*^p53KO^*cells (Figure S4A-B). Thus, to optimize our screen for evaluation of cell death, we increased the dose to induce substantial lethality, and shortened the assay time, such that evolution of the drug-treated sgRNA population would be driven primarily by variations in the degree of cell death between clones, rather than by variations in the degree of proliferation. Treatment with 5 µM etoposide for 4 days resulted in ∼50% lethal fraction in both cell types, without causing a population “bottleneck”, which would deteriorate assay sensitivity (Figure 3B-C). Preliminary analyses, such as the high correlation among replicates and degree of dropout among essential genes, suggest that data quality was not compromised by the atypical assay conditions used in our screen (Figure 3D and S4C-E).

**Figure 3:**
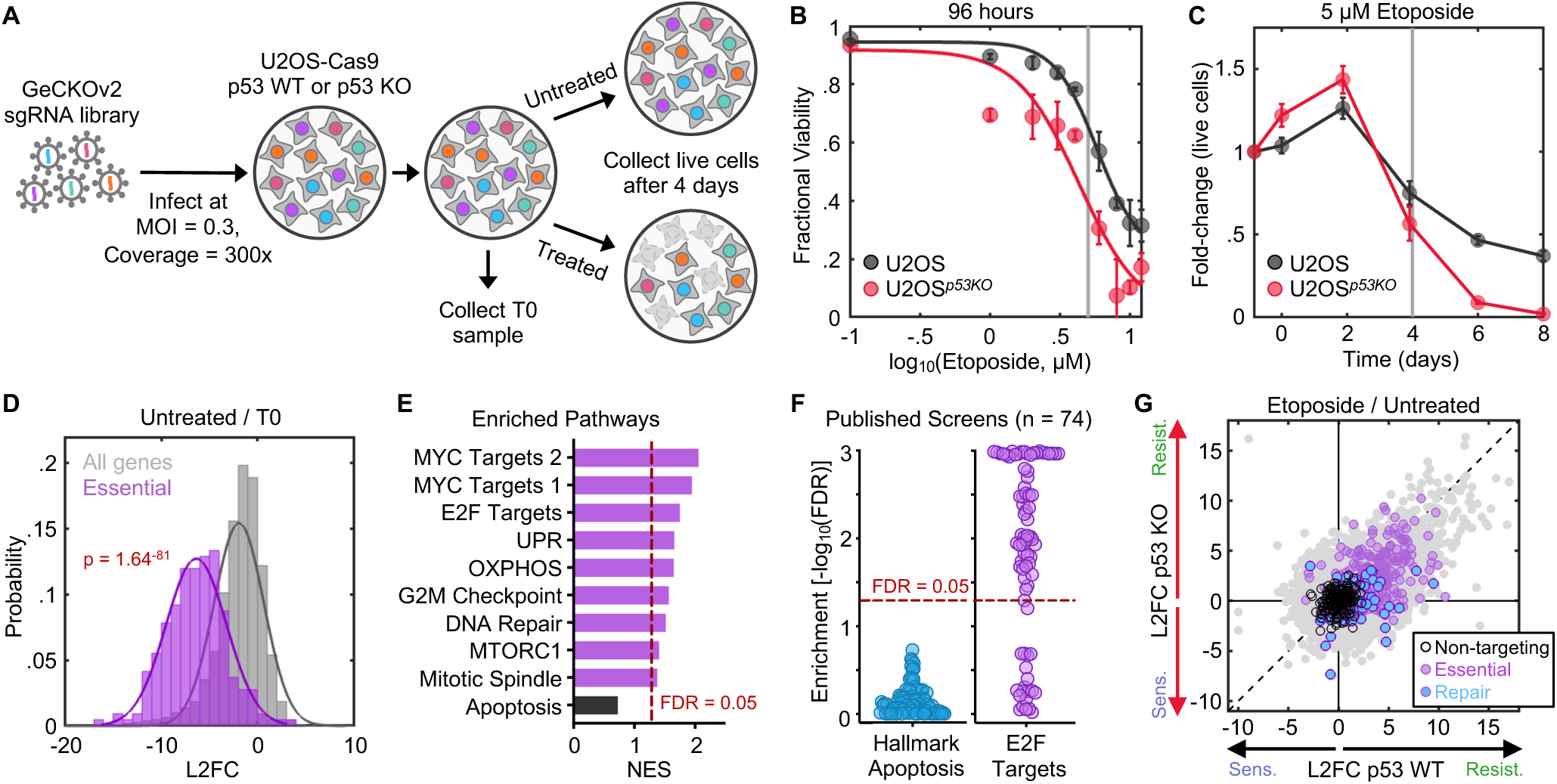
Genetic screens fail to identify death regulatory genes due to confounding effects caused by varied growth rates. **(A)** Schematic of pooled screen **(B-C)** Parameterization of drug dose and assay time for pooled screen. Screen dose (B) and assay length (C) were selected to produce intermediate levels of lethality while maintaining a population size large enough for > 300× coverage of the sgRNA library throughout the assay. **(D)** Distribution of all genes compared to core essential genes in the untreated vs. T0 sample for U2OS cells. KS test p-value shown. **(E)** GSEA for U2OS etoposide vs. untreated samples, showing most enriched gene signatures. Apoptosis is not significant, shown for comparison. **(F)** GSEA-based analysis for 74 published genome-wide screens of apoptotic agents. Apoptotic genes are consistently missed, while screens typically enrich for known proliferation genes. **(G)** Gene-level log_2_ fold change (L2FC) for U2OS (WT) compared to U2OS*^p53KO^* (KO). DNA repair genes and core essential genes shown to demonstrate enrichment for genes that reduce growth fitness in chemo-genetic profiling data. See also Figure S3 and S4.

Functional genetic screens are typically analyzed by computing guide-level or gene-level fitness scores, which are generated by comparing the relative abundance of guides or genes between the drug-treated and untreated conditions (*e.g.,* log2-fold change, L2FC)^44–46^. Notably, the comparison of “drug-treated vs. untreated” is precisely the same as “relative viability”, which fails to accurately capture the contribution of cell death. Thus, we reasoned that conventional analysis methods would also fail to identify death regulatory genes. Indeed, analysis of our data using conventional L2FC methods, and re-analysis of 74 published functional genetic screens of apoptotic drugs, did not identify apoptotic regulatory genes (Figure 3E-F). Conventional analyses were, however, effective at identifying genes that regulate proliferation (Figure 3E-F). Further inspection also revealed clear evidence of confounded insights due to growth rate variation among knockout clones. For example, knockout of DNA repair genes should phenocopy higher doses of etoposide, and thus should sensitize cells to etoposide (*e.g.,* negative L2FC). However, DNA repair genes generally scored with increased L2FC, commonly interpreted as promoting drug resistance (Figure 3G). A similar phenotype is observed for essential genes, which drop out over time due to low growth rates (Figure 3G)^47^. Thus, as with *in vitro* drug response analysis, conventional analysis methods used in functional genetic screening fail to identify death regulatory genes due to the confounding effects of variations in growth rates between knockout clones.

The effect of a gene knockout on cell death would be observable if comparing the relative abundance of KO compared to WT clones (Δ2, Figure 4A). However, in pooled genetic screens the effect of a gene knockout is scored across treatment conditions (KO_tx_ vs KO_unt_), using WT as a normalizing control (Figure 4A). Thus, a conventional gene effect score is only partially influenced by the death rate (Figure 4B). Furthermore, using this score to infer the death regulatory function of a gene results in a range of possible errors (Figure 4C), including an inverted inference if the growth rate defect is strong enough, as is seen with DNA repair genes (Type III error, Figure 4C-D). Thus, accurately identifying death regulatory genes from chemo-genetic profiling data requires a calculation of the drug-induced death rate, rather than the relative population size or relative growth fitness.

**Figure 4:**
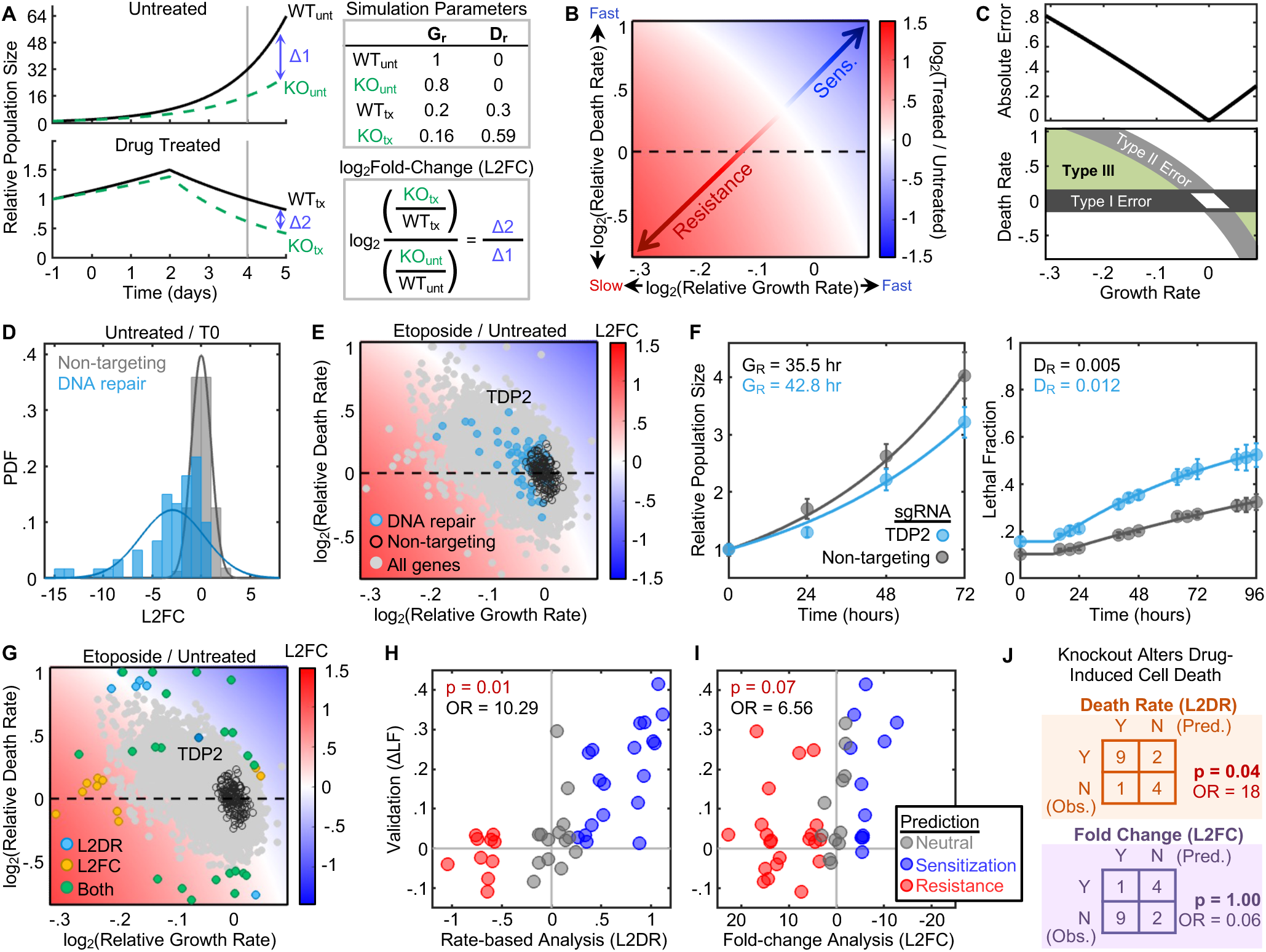
Death rate-based analysis accurately identifies genes that regulate drug-induced death. **(A)** Simulation to highlight conceptual issues with common analysis methods. Pooled genetic screens do not score the relationship between WT and KO cells, but instead, the relative abundance of KO cells in treated and untreated populations (L2FC). In the example, KO cells die twice as fast, but this is obscured by their modest 20% growth defect, which creates a large difference in population size in the fast-growing untreated population. In the example, L2FC is positive in spite of a higher death rate in the knockout cells. Positive L2FC (*i.e.*, enrichment) is generally interpreted as drug resistance **(B)** Full simulation for all combinations of growth rates and death rates. Phase diagram shows how changes to growth/death combine to create population sizes that are commonly interpreted as drug sensitization (Sens.) or drug resistance. **(C)** Error in death rate inferences from L2FC values depending on the growth rate of knockout clones. Absolute error as a function of growth rate (*top*). Phase diagram showing type of error as a function of growth rate and death rate (*bottom*). **(D)** Probability density function (PDF) for non-targeting sgRNAs or DNA repair genes in Untreated vs. T0 comparison. Knockout of DNA repair genes causes reduced growth rate. **(E)** Gene-level chemo-genetic profiling data for etoposide vs. untreated in U2OS cells, projected into phase diagram. DNA repair genes and non-targeting sgRNAs highlighted. **(F-J)** Validation of screen. **(F)** Example validation of TDP2, a DNA repair gene. **(G)** Expanded validation of 40 genes that score strongly by death rate or fold change analysis. **(H)** Validation results for 40 genes compared to predictions based on the death rate-based analysis (log_2_ death rate, L2DR). **(I)** As in panel H, but compared to L2FC. For panels H and I, odds ratio (OR) and p-values shown based on one-tailed Fisher’s exact test (*i.e.*, hypergeometic distribution). **(J)** Fisher’s exact test for the subset of gene knockouts within the validation set with reduced growth rates. See also Figure S5.

For *in vitro* drug response analyses, accurate calculation of the death rate requires counting both the live and dead cell fractions^27,33^. While recovery of dead cells is possible in the context of apoptotic death due to the relative stability of the apoptotic corpse^48,49^, recovering intact dead cells may not be possible in the context of non-apoptotic death due to cell rupture and instability of necrotic cells. Thus, we devised a new strategy for inferring the death rate from the data that is typically available in chemo-genetic profiling studies (*i.e.*, sequencing counts proportional to the number of live cells over time in treated and untreated conditions). Our general strategy was to infer the drug-induced death rate from a combination of 3 experimental insights: 1) the relative abundance of each sgRNA in treated versus untreated groups (*i.e.,* the conventional gene effect score), 2) the observed growth rate defect for each sgRNA in untreated vs. T0 comparison, and 3) the experimentally observed coordination between growth and death from GRADE-based analysis (see Methods and Figure S5A-C). In the context of the high levels of DNA damage screened in our assay, death only occurs in growth arrested cells, which simplifies the inference of death rate by limiting the possible ways that perturbations to growth rate and death rate integrate to create a given number of cells (Figure S5A)^27^. Using this inference approach, we computed the gene-level growth rates and death rates for each single gene knockout (Figure 4E).

Our analysis confirmed that knocking out DNA repair genes increases the death rate of cells exposed to DNA damage (*e.g.*, positive values on the y-axis), but that this phenotype is not apparent due to the slow growth rate of these cells in the untreated condition (Figure 4E). We also identified “false negatives” (*i.e.*, Type II error) that could be rescued by our death rate analysis. For instance, TDP2 is a DNA repair protein that specifically repairs adducts created by etoposide, and knocking out TDP2 increases etoposide potency^43,50^. At the doses and times used in our chemo-genetic profiling, a conventional L2FC-based analysis scores TDP2 at approximately 0, erroneously suggesting that knocking out TDP2 has no effect on etoposide sensitivity. Our death rate analysis reveals that the neutral L2FC results from a combination of increased death rate in the drug-treated condition and decreased growth rate in the untreated condition. To validate these model-inferred predictions, we tested TDP2-targeting sgRNAs for their effect on both cell proliferation and cell death, finding, as predicted, that knockout of TDP2 resulted in a slower growth rate in the absence of exogenous DNA damage, and a faster death rate in the presence of etoposide (Figure 4F).

Our method for inferring the drug-induced death rate relies on a critical assumption, that cell death occurs following DNA damage in a non-growing population. While this assumption is experimentally validated for wild-type cells, it may not always be true in the context of gene deletions which could alter the coordination of growth and death. Thus, we performed a large-scale validation of our death rate inferences for different classes of predictions (Figure 4G and S5D-E). We focused on the top 20 genes predicted to increase or decrease the death rate using our inference approach. For comparison, we also evaluated the top 20 genes predicted to cause drug sensitization/resistance based on a conventional L2FC-based analysis. Overall, death rate-based analysis of our chemo-genetic profiling data was strongly predictive of the death regulatory function of each gene (Figure 4H).

For the traditional analysis methods, our chemo-genetic profiling data was not significant for predicting the cell death regulatory function of the 40 tested genes (Figure 4I). The data, however, were also clearly non-random (OR = 6.6) and may have been significant if we tested a larger set of genes. We note, however, that these two analysis methods only produce conflicting predictions for gene knockouts that cause reduced growth rates. Thus, to directly compare rate-based and population fold-change-based analyses, we focused on the subset of genes whose deletion caused a reduced growth rate. Regardless of the growth rate observed for each single gene knockout, our death rate-based analysis continued to accurately predict the death regulatory function of each gene (OR = 18; Figure 4J and S5F). Importantly, for gene knockouts with a reduced growth rate, a conventional fold-change analysis produced predictions of the cell death regulatory function that were approximately twenty times worse than random predictions (OR = 0.06; Figure 4J and S5F). Taken together, these data reveal that commonly used analysis methods are insensitive and inaccurate for studying cell death, and that a death rate-based analysis can improve the accuracy and interpretation of the death regulatory function of each gene.

### In the absence of p53 DNA damage activates a mitochondrial respiration-dependent form of necrosis

We next sought to determine if our death rate data were sufficient for interpreting mechanisms of drug-induced cell death, and if these data could be used to determine which type of cell death is activated by DNA damage in the absence of functional p53. Our chemo-genetic profiles identified 502 genes that modulate the etoposide-induced death rate when deleted in wild-type U2OS, and 755 genes that modulate the etoposide-induced death rate when deleted in U2OS*^p53KO^*. Importantly, the overwhelming majority of these “hits” were observed in only one of the two genetic contexts (*e.g.*, 619 of 755 genes were unique to U2OS*^p53KO^* cells); and overall, we observed a poor correlation between the death rates in the presence and absence of p53 (Figure 5A). This is in stark contrast to what we observed with a conventional fold change-based analysis, in which scores were similar for p53 WT and KO cells (Figure 3G). Additionally, many expected phenotypes were observed with the correct directionality in our death rate analysis. For instance, deletion of p53, or critical p53-target genes such as p21 (CDKN1A), increased the death rate exclusively in the p53 WT background, whereas deletion of critical DNA repair factors such as TDP2 increased the death rate in both backgrounds (Figure 5A).

**Figure 5:**
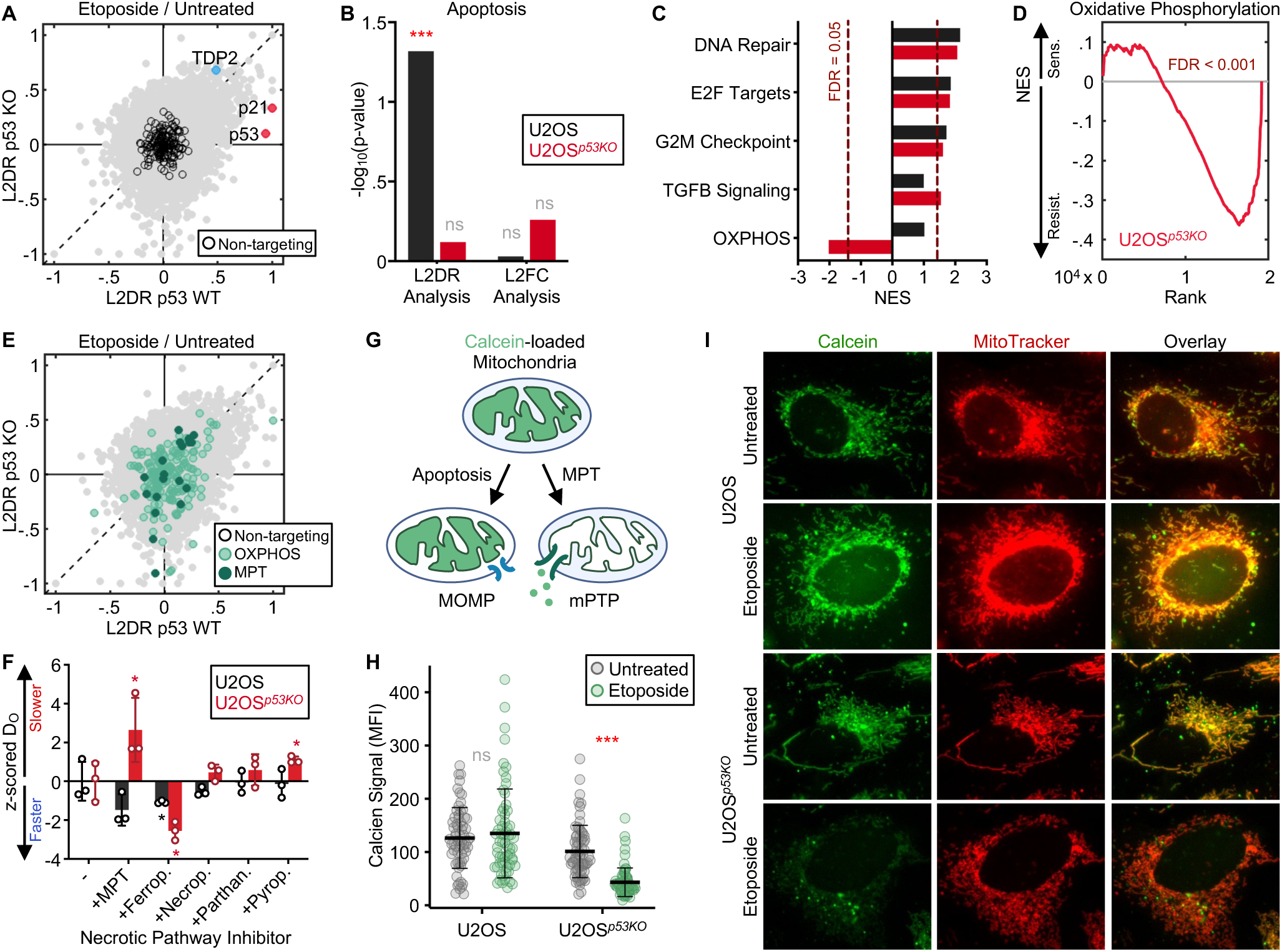
DNA damage activates a respiration-dependent form of necrotic death in the absence of p53. **(A)** Gene-level log_2_ death rate (L2DR) for U2OS (WT) compared to U2OS*^p53KO^* (KO). TDP2, p53, and p21 are highlighted to demonstrate directionality of known controls. **(B)** Pathway-level enrichment (GSEA) of apoptotic genes from U2OS and U2OS*^p53KO^* chemo-genetic screens analyzed with a fold-change or rate-based analysis. Apoptotic genes are enriched only in U2OS, analyzed using L2DR. **(C)** GSEA for the death rate of etoposide treated cells, showing signatures most enriched in U2OS*^p53KO^*. Negative normalized enrichment scores (NES) indicate a decrease in death rate, positive NES indicates an increase in death rate. **(D)** Running enrichment score for the oxidative phosphorylation (OXPHOS) signature in U2OS*^p53KO^*. **(E)** L2DR for U2OS and U2OS*^p53KO^* cells, highlighting OXPHOS genes and known regulators of MPT-driven necrosis. **(F)** U2OS and U2OS*^p53KO^* treated with 31.6 µM etoposide, zVAD, and the indicated death pathway inhibitor for 48 hours. The death onset time (D_O_) of each was z-scored relative to the effect of etoposide and zVAD alone (*left-most group*). **(G-I)** Calcein-cobalt assay to evaluate opening of the mPTP. **(G)** Schematic of how the cobalt-calcein assay differentiates apoptosis and MPT. **(H)** Measurement of calcein signal in U2OS and U2OS*^p53KO^* treated with 31.6 µM etoposide for 36 hours. >50 individual cells scored in each condition. **(I)** Representative images of treated and untreated cells. MitoTracker Red was used to mask and quantify calcein fluorescence. See also Figure S6.

To determine if these data are sufficient for interpreting the mechanism of cell death activated by etoposide, we focused first on U2OS cells, where death occurs via canonical apoptosis. Importantly, in U2OS, gene deletions that alter the death rate were significantly enriched for known apoptotic regulatory genes (Figure 5B). These effects were not observed in the U2OS*^p53KO^* background. Furthermore, the distinction between p53-proficient and p53-deficient cells was not observed when we used a conventional fold change-based analysis (Figure 5B). These data further confirm a lack of DNA damage-induced apoptosis in p53 KO cells, and the unique sensitivity of our death rate-based analysis method for revealing the mechanism of cell death.

Most non-apoptotic death subtypes remain poorly annotated, but in general, for each death subtype at least some unique effector enzymes are known^3^. To determine the type of cell death activated by etoposide in the absence of p53, we used our death rate analyses to identify genes/genetic signatures that specifically modulate the etoposide-induced death rate in U2OS*^p53KO^* cells, and not in the etoposide-treated U2OS cells. This analysis revealed a unique dependency in U2OS*^p53KO^*cells for genes involved in oxidative phosphorylation (OXPHOS) (Figure 5C-D). Deletion of OXPHOS regulatory genes rescued viability in etoposide-treated U2OS*^p53KO^*; however, knocking out these genes had no effect on the death rate in U2OS cells (Figure 5E and S6A-B). Critically, if these data were analyzed using a conventional fold-change analysis method, OXPHOS would score as promoting “resistance” in both p53-proficient and – deficient settings (Figure 3E). However, our death rate analysis clarifies that, while OXPHOS genes have different effects on death in the presence and absence of p53, these genes have similar effects on growth in both contexts. Loss of OXPHOS inhibits growth in both cell types, and this shared dependency drives the specious drug resistance phenotype that is observed in a conventional analysis (Figure S6A-B). Thus, the critical differences in mechanisms that regulate lethality in the presence and absence of p53 would have been missed using conventional analyses.

Some OXPHOS regulatory proteins are components of a mitochondrial respiration-dependent form of cell death called mitochondrial permeability transition (MPT)-driven necrosis^3,51^. MPT-driven necrosis is thought to be caused by opening of a mitochondrial permeability transition pore (mPTP), loss of mitochondrial inner membrane integrity, and mitochondrial rupture. These events ultimately cause a morphologically necrotic death. Many of the molecular and mechanistic details of MPT-driven necrosis remain unknown and/or are controversial; however, it is well-established that MPT-driven necrosis depends on the activity of cyclophilin D (CYPD; MPT-driven necrosis is also known as CYPD-dependent necrosis)^52^. CYPD-dependent death is generally evaluated using the cyclophilin family inhibitor, cyclosporin A (CsA)^51,53^. In our initial evaluation of non-apoptotic death pathways, we did not evaluate MPT-driven necrosis, as CsA itself activated apoptotic death in U2OS cells (Figure S6C). To more carefully evaluate whether MPT-driven necrosis contributes to DNA damage-induced cell death in the absence of p53, we tested CsA in combination with the caspase inhibitor zVAD. As expected, zVAD neutralized the lethality of CsA (Figure S6D). In the presence of zVAD, CsA slowed the onset of etoposide-induced death in U2OS*^p53KO^*, but did not affect the etoposide response in U2OS (Figure 5F). Additionally, we tested inhibitors of other necrotic pathways for which mitochondrial activity contributes to lethality. None of these inhibited death in U2OS*^p53KO^*cells in the presence or absence of zVAD (Figure 5F and S3A-B).

To further evaluate whether MPT-driven necrosis is activated by DNA damage in the absence of p53, we sought to monitor opening of the mPTP. We used the Co2+-calcein assay, in which cytoplasmic fluorescence of calcein is quenched by cobalt; opening of the mPTP facilitates cobalt entry and quenching of mitochondrial calcein fluorescence^54^. In U2OS*^p53KO^* cells, etoposide exposure caused a significant decrease in mitochondrial calcein fluorescence (Figure 5G-I). Similar results were found in other p53-deficient cell lines, with mitochondrial calcein fluorescence being lost following exposure to a lethal dose of etoposide (Figure S6E). In U2OS and other p53 WT cells, we observed no change in mitochondrial calcein fluorescence following exposure to etoposide (Figure 5G-I and S6E). Taken together, these data demonstrate that DNA damage activates MPT-driven necrosis in the absence of p53.

### High basal respiration due to mitochondrial supercomplex formation primes cells for MPT-driven necrosis

We next aimed to identify features that drive the propensity to activate MPT-driven necrosis in p53-deficient cells. We had originally suspected MPT-driven necrosis based on the differential genetic dependencies that we observed for genes that regulate OXPHOS. To investigate this issue further, we measured the cellular oxygen consumption rate (OCR) in U2OS and U2OS*^p53KO^*, using the Seahorse Extracellular Flux assay. While drug exposure induced a modest decrease in the spare respiratory capacity of both cell types, we observed, independent of drug therapy, that U2OS and U2OS*^p53KO^* have strikingly different respiratory profiles (Figure 6A). Compared to U2OS, U2OS*^p53KO^* cells have increased essentially all aspects of respiration, including the basal OCR and spare respiratory capacity (Figure 6B).

**Figure 6:**
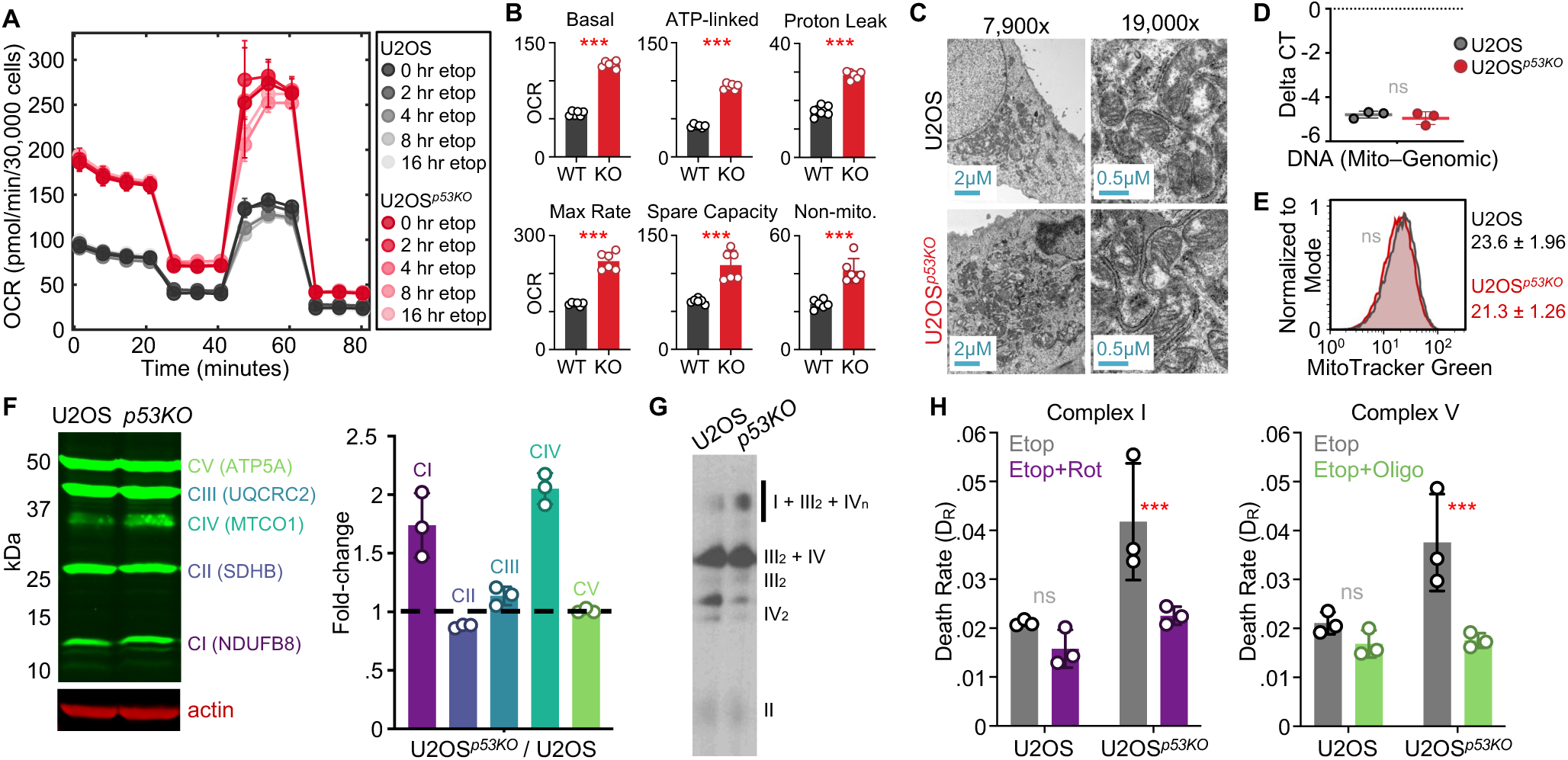
High basal respiration due to mitochondrial supercomplex formation primes cells for MPT-driven necrosis. **(A-B)** Oxygen consumption rate (OCR) in U2OS and U2OS*^p53KO^* treated with etoposide for indicated times. **(A)** Full mitochondrial stress test response profile. **(B)** Respiration rates for WT and KO cells. **(C)** TEM of U2OS and U2OS*^p53KO^*. **(D-E)** Mitochondrial abundance in U2OS and U2OS*^p53KO^* evaluated using (D) qPCR of mitochondrial DNA or (E) MitoTracker Green fluorescence level. **(F)** Relative abundance of ETC complexes I-V. (*left*) representative immunoblot. (*right*) Quantification of three biological replicates. **(G)** Blue native PAGE of ETC complexes. Representative blot shown from three biological replicates. **(H)** Death rate following etoposide exposure (Etop) in the presence of Rotenone (Rot) or Oligomycin (Oligo). For all panels with error bars, data are mean ± SD from 3 experimental replicates.

A possible explanation for the observed difference in OCR could be that U2OS*^p53KO^* have increased numbers of mitochondria. Indeed, p53 has been demonstrated to regulate mitochondrial fission-fusion dynamics^55^. However, we did not observe any differences in mitochondrial morphology or abundance between U2OS and U2OS*^p53KO^*(Figure 6C-E). An alternative mechanism by which OCR could increase is through increased respiratory efficiency in U2OS*^p53KO^*. Many distinct mechanisms control mitochondrial efficiency, often through regulation of the electron transport chain complexes, Complex I – V^56^. To evaluate this further, we quantified the relative abundance of five ETC complex proteins. We observed increased levels of proteins in Complex I, Complex III, and Complex IV (Figure 6F). ETC Complexes I, III, and IV can form a single macromolecular complex, called a supercomplex (SC)^57,58^, which has been shown to increase respiratory efficiency^59^. Therefore, we determined if SC formation is enhanced in U2OS*^p53KO^* cells. Using blue native PAGE (BN-PAGE), we found in U2OS*^p53KO^* cells that the monomeric bands for constituents of mitochondrial SCs, such as Complex IV, are depleted, while high molecular weight bands that are consistent with the expected mass of SC are enriched relative to U2OS (Figure 6G).

Next, we aimed to determine if SC formation and/or increased respiration are required for the activation of MPT-driven necrosis in p53-deficient cells exposed to etoposide. We were unable to perturb SC formation without also reducing mitochondrial respiration. Nonetheless, to directly evaluate if high respiration contributes to the propensity to activate MPT-driven necrosis, we measured etoposide sensitivity in the presence or absence of rotenone, to inhibit ETC Complex I, or oligomycin, to inhibit ETC Complex V. For both inhibitors, we observed no effect in the p53-proficient U2OS background; however, both rotenone and oligomycin inhibited cell death in p53-deficient U2OS*^p53KO^* (Figure 6H). Taken together these data demonstrate that the high basal respiratory levels in these p53-deficient cells prime cells for MPT-driven necrosis.

### Metabolic vulnerabilities unique to p53-deficient cells can be targeted to potentiate DNA damage sensitivity

Having found that the distinct metabolic states of p53-proficient and p53-deficient cells contribute to the activation of different forms of cell death, we next aimed to identify strategies for potentiating the activation of non-apoptotic cell death in p53-deficient cancer cells. To determine ways to achieve this, we used mass spectrometry-based metabolite profiling to identify targetable differences in the metabolic states of U2OS and U2OS*^p53KO^*, both at steady-state and following etoposide exposure. Overall, we quantified the abundance of 463 metabolites. The basal steady-state metabolite profiles were very similar for these isogenic cells, with significant differences observed for only 11 metabolites (Figure S7A). More pronounced differences were observed following etoposide exposure, with 118 metabolites significantly enriched or depleted following drug exposure, in U2OS and/or U2OS*^p53KO^*cells (Figure 7A). To identify opportunities for specifically targeting p53-deficient cells, we identified metabolic processes that were altered by etoposide only in the absence of p53. The metabolites altered following etoposide exposure were enriched within 10 total metabolic pathways; only 3 closely related pathways were altered uniquely in the U2OS*^p53KO^*background: Glycolysis, Pentose Phosphate Pathway (PPP), and TCA Cycle (Figure 7B). In general, observed metabolites within these 3 pathways were depleted in U2OS*^p53KO^*cells following drug exposure; similar changes were not observed in p53-proficient U2OS cells (Figure 7C and S7B).

**Figure 7:**
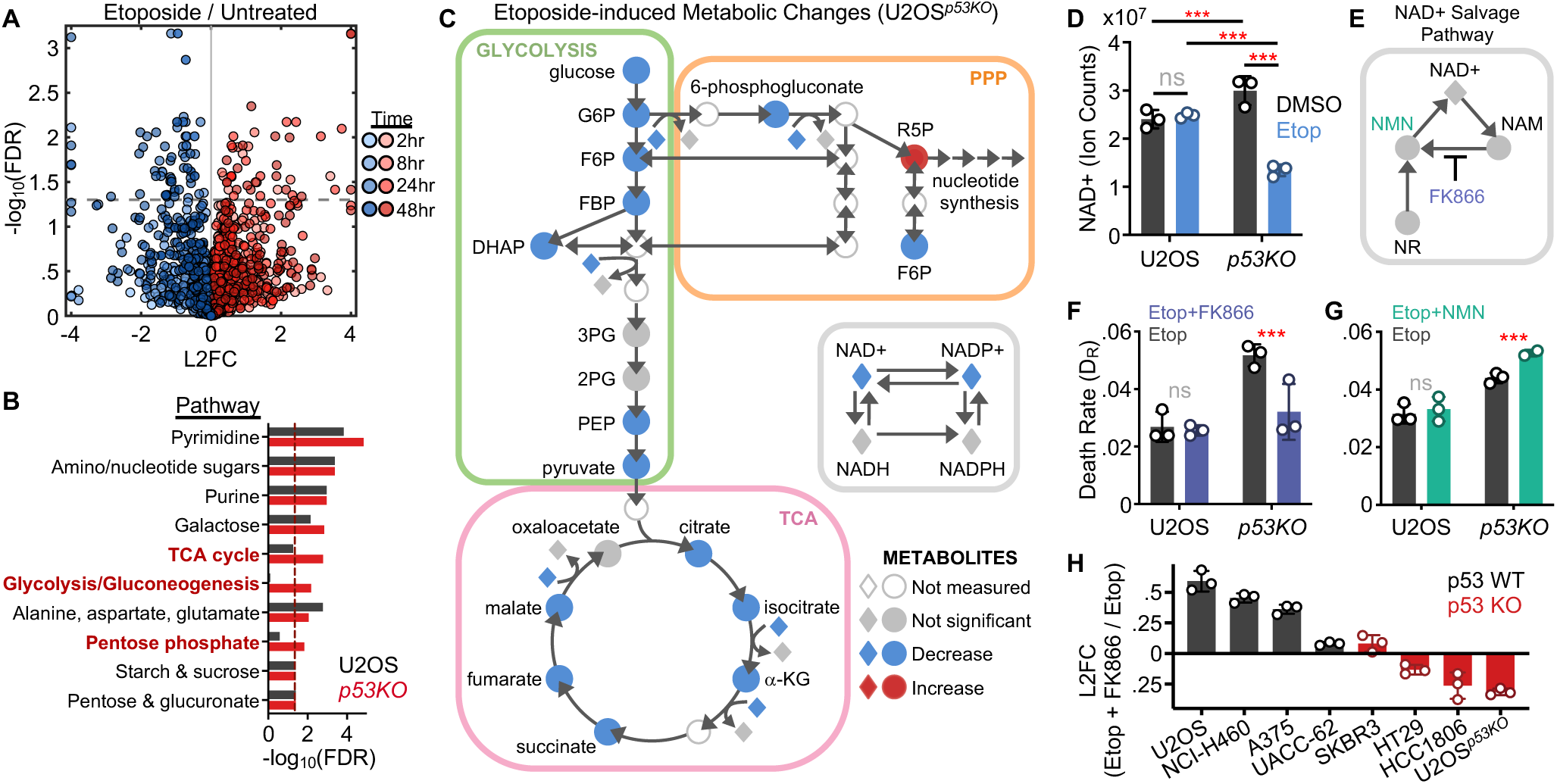
Metabolic vulnerabilities unique to p53-deficient cells can be targeted to potentiate DNA damage sensitivity. **(A)** LC-MS based profiling of metabolites. Volcano plot shows log2 fold change (L2FC) in metabolite levels in p53 KO following etoposide exposure for indicated times. **(B-C)** Pathway enrichment among significantly altered metabolites. **(B)** Highlighted in red are pathways that are changed in p53 KO but not p53 WT using metabolic pathways defined by MetaboAnalyst. **(C)** Detailed view of metabolites changed within glycolysis, PPP, and TCA pathways. Metabolites that differ following 24 or 48 hour etoposide exposure are highlighted red/blue. **(D)** NAD+ levels at steady-state in treated or untreated samples. **(E-H)** Perturbation of NAD+ levels. **(E)** Schematic of the NAD+ Salvage Pathway. **(F)** Death rate (D_R_) measured using the FLICK assay 48 hours after drug exposure in the presence or absence of 1 µM FK866. **(G)** As in panel (F) but in the presence or absence of 1.58 mM NMN. **(H)** FK866 sensitivity across a panel of cell lines with functional (grey) or dysfunctional (red) p53. Abbreviations: glucose 6-phosphate (G6P), fructose 6-phosphate (F6P), fructose 1,6– bisphosphate (FBP), dihydroxyacetone phosphate (DHAP), 3-phosphoglycerate (3PG), 2– phosphoglycerate (2PG), phosphoenolpyruvate (PEP), ribose 5-phosphate (R5P), a-ketoglutarate (a-KG), nicotinamide (NAM), nicotinamide mononucleotide (NMN), nicotinamide riboside (NR), uridine monophosphate (UMP). For all panels with error bars, data are mean ± SD from 3 experimental replicates. See also Figure S7 and S8.

Decreased abundance of glycolytic, PPP, and/or TCA metabolites following drug exposure could have occurred due to DNA damage-induced changes in the use of these metabolic pathways. Alternatively, these changes may have resulted from “survivor bias”, caused by preferential death of cells with high levels of these metabolites. To distinguish between these possibilities, we began by using isotope tracing with labeled glucose (Figure S8A). The degree to which a given intermediate metabolite within the glycolytic, PPP, or TCA pathway is derived from the activity of one of these pathways can be represented as the ratio of the labelled downstream metabolite relative to its labelled upstream precursor (*e.g.,* fumerate+2/citrate+2)^60^. Based on this analysis, etoposide did not induce significant changes in the use of glycolysis (Figure S8B). Etoposide altered the use of PPP and TCA pathways; however, the observed changes were nearly identical in U2OS and U2OS*^p53KO^*cells (Figure S8B). Thus, for the depleted metabolites within glycolysis, PPP, and TCA pathways, the specific loss of metabolites in the absence of p53 did not result from differences in the use of these pathways within these cells.

Based on these data, we sought to directly test whether high levels of metabolites within TCA, PPP, and/or glycolysis pathways sensitize p53-deficient cells to DNA damage. Although very few metabolites differed in untreated conditions between U2OS and U2OS*^p53KO^* cells, we observed a striking difference for NAD+. NAD+ is a cofactor involved in multiple metabolic reactions within the TCA, PPP, and glycolysis pathways (Figure 7C). Prior to etoposide exposure, we observed that NAD+ is higher in U2OS*^p53KO^*compared to U2OS cells (Figure 7D). At long time points (*i.e.,* longer than the death onset time) following etoposide exposure, however, NAD+ is significantly decreased in U2OS*^p53KO^*, but remains unchanged in U2OS cells (Figure 7D). This pattern of changes is consistent with survivor bias, in which high levels of NAD+ sensitize cells to DNA damage, only in the p53-deficient context. To directly test this model, we exposed cells to the NAMPT inhibitor, FK866. NAMPT is the rate-limiting enzyme in the NAD salvage pathway, which promotes production of NAD+ (Figure 7E)^61,62^. We observed that FK866 inhibits the lethality of etoposide, and this phenotype is observed only in p53-deficient cells (Figure 7F). To further test this model, we also tested supplementation with NMN, the precursor metabolite used in generation of NAD+ through the salvage pathway^63^. As predicted, we observed that NMN supplementation enhances the lethality of etoposide, and again, this phenotype is observed only in the p53-deficient context (Figure 7G). Finally, to evaluate the generalizability of our findings, we tested the lethality of etoposide in the presence and absence of FK866, in a panel of genetically diverse p53-proficient and p53-deficient cancer cells. These data confirm our general finding that NAD+ promotes the lethality of etoposide only in p53-deficient cancer cells (Figure 7H). Taken together, these findings reveal a unique metabolic vulnerability which can be exploited to potentiate the lethality of DNA damage, specifically in the context of p53-deficiency.

## DISCUSSION

In this study, we describe a previously uncharacterized function for p53 in biasing how cells respond to lethal levels of DNA damage. It has long been known that p53 is required for robust activation of apoptosis following DNA damage. However, prior studies have generally concluded that loss of p53 promotes survival of cells treated with DNA damaging agents. In contrast, in cancer cells that are naturally deficient for p53, and in cells genetically engineered to lack p53, we find that loss of p53 does not decrease the levels of DNA damage-induced cell death. Instead, cells lacking p53 simply die using a non-apoptotic form of death, MPT-driven necrosis. Furthermore, using metabolomic profiling, we identified that increasing levels of NAD+ can enhance the lethality of DNA damage, specifically in p53-deficient cancer cells.

While prior studies have found that p53-deficient cells can use alternate pathways to activate apoptotic cell death, such as a caspase-2-dependent signaling pathway that is inhibited by Chk1^64^, non-apoptotic forms of cell death have not been well studied in these cells. Our prior studies have revealed a phenomenon that is commonly observed in the context of cell death called “single agent dominance”, in which faster acting death pathways suppress the activation of slower acting pathways^9^. Thus, while other forms of death were not observed in the panel of p53-deficient cell types used in this study, this potential variation highlights that many different types of death may be possible once the canonical cell intrinsic apoptotic response is no longer a dominant mechanism. Our study suggests that MPT-driven necrosis is the dominant form of cell death in DNA damaged cells that lack functional p53. Furthermore, as we highlighted in this study, the non-apoptotic form of death that we found would likely not have been identified in prior studies due to the common use of assays that are biased towards proliferative effects.

Importantly, use of proliferation-focused assays, or measurements such as RV, cannot fully explain why previous studies failed to observe robust activation of DNA damage-induced non-apoptotic death. Increased relative viability is often interpreted as increased cell survival, particularly when increased RV is also associated with loss of observable dead cells. Thus, a second issue that clearly contributes to under-scoring non-apoptotic death is the historical bias towards evaluating cell death using markers that are specific to apoptotic bodies – caspase-3 cleavage, Annexin V positivity, chromosome condensation, and membrane blebbing, etc. – and these features are absent from dead cells following DNA damage-induced non-apoptotic cell death. In this study, our evaluations of cell death were made using the FLICK assay, a plate reader-based assay that produces a death-specific measurement. Importantly, FLICK requires only the loss of membrane integrity, and these measurements are obtained without requiring the collection, handling, or separation of dead cells, which may bias against cells that rupture following necrotic death^9,26^. These findings highlight that studies of anti-cancer drug responses should include either direct measurements of each relevant form of cell death, or death-specific measurements that are agnostic to the type of death, with the precise death subtype inferred through genetic and chemical dependencies, as in this study.

Additionally, a surprising observation from this study is our finding that pooled loss of function genetic screens are systematically biased against interpreting the death regulatory function of genes. Intuitively, growth fitness (*i.e.,* the relative population growth rate) is influenced by changes to both the cell proliferation rate and the death rate. Thus, functional genetic screens should, in principle, be sensitive to knockout of death regulatory genes. Empirically, we observe that previously published screens routinely fail to identify essential apoptotic regulators, even when the death in question is known to be apoptotic. Likewise, for the data collected in this study, conventional analysis methods produced erroneous insights, and these methods failed to identify the genetic dependencies for DNA damage-induced lethality, or how these dependencies vary in the presence and absence of p53. The critical insights generated in this study, such as the varied use of OXPHOS in p53-proficient and –deficient cancer cells, would have been missed using conventional analysis approaches. The modeling and analysis in this study reveals three culprits for the common lack of sensitivity to death regulatory genes that has been observed in functional genomic studies: use of non-lethal drug concentrations, use of long assay lengths that bias towards proliferative phenotypes, and use of analytical methods that fail to address the confounding effects of growth rate variation in the untreated samples. This latter issue is problematic because knocking out a death regulatory gene also tends to affect the cell proliferation rate.

Growth rate variation has similarly been recognized as a confounding influence in the evaluation of drug sensitivity. For *in vitro* drug sensitivity, the drug-induced death rate can be accurately scored by measuring both the live and dead cell populations. Because it is often not possible to make these measurements for non-apoptotic death, we generated a new method for inferring the drug-induced death rate from a combination of three insights: 1) the population size at assay end point, 2) the experimentally measured growth rate for each clone in the pool, and 3) the coordination of growth and death from GRADE-based analysis. While the model we used for our analysis was specific for the growth-death coordination that is observed for DNA damage, with appropriate modifications to this model, these methods should be usable in any other drug context. Moreover, while other methods, such as direct sequencing of dead cells following sorting, can be effective in the context of apoptotic death, the methods used in this study are likely to be the best methods for identifying the genetic dependencies for non-apoptotic types of death.

Taken together, our findings also reveal that the methods used throughout the community for evaluating drug sensitivity have generated a systematic bias, limiting our understanding of non-apoptotic forms of cell death. In the context of p53-deficient cancers, we found unexpectedly high levels of non-apoptotic death following DNA damage. Because necrotic cells are difficult to recover and because these cells lack the conventional hallmarks of apoptotic cells, non-apoptotic death can go undetected, and these phenotypes would previously have been erroneously attributed to lack of proliferation. As we report herein, the genetic dependencies for apoptotic and non-apoptotic forms of cell death are distinct, even when death is activated by a common stimulus, like etoposide-induced DNA damage. Thus, efforts to personalize lethal drug therapies will likely benefit from understanding the shared versus death pathway-specific genetic dependencies for commonly used therapeutic agents. The extent to which targeting non-apoptotic death pathways in p53-deficient cancers would improve therapeutic responses remains to be determined, but our studies reveal the prominent use of MPT-driven necrosis in p53-deficient cancers, and the genetic determinants for DNA damage-induced and MPT-driven cell death.

## METHODS

### Cell lines and reagents

A431, A549, BT549, HCC1143, HCC1806, HT-29, MCF7, MCF10A, MDA-MB-231, MDA-MB-436, MDA-MB-453, MDA-MB-468, SK-BR-3, and U2OS cells were obtained from the American Type Culture Collection (ATCC). A375, HT-1080, Malme-3M, NCI-H460, and UACC62 cells were acquired from the Green lab (UMass Chan Medical School). Caki-1 cells were a acquired from the Kim lab (UMass Chan Medical School). A375, A431, A549, MCF7, MDA-MB-231, MDA-MB-436, MDA-MB-453, MDA-MB-468, and U2OS cells were grown in DMEM (Corning, #10-017-CV). HT-1080 cells were grown in EMEM (ATCC, #30-2003). Malme-3M cells were grown in IMDM (Thermo Fisher Scientific, #12440053). Caki-1, HT-29, and SK-BR-3 cells were grown in McCoy’s 5A medium (Corning, #10-050-CV). BT549, HCC1143, HCC1806, NCI-H460, and UACC62 cells were grown in RPMI 1640 medium (Corning, #10-040-CV). MCF10A cells were grown in DMEM-F12 medium (Thermo Fisher Scientific, #11320033) supplemented with 5% horse serum, 20 ng/ml epidermal growth factor (EGF), 0.5 mg/ml hydrocortisone, 100 ng/ml cholera toxin (Sigma-Aldrich, #C8052-2MG), 10 μg/ml insulin (Life Technologies, #12585014), and penicillin-streptomycin (Corning, #30-002-CI). Each media (except DMEM-F12) was supplemented with 10% fetal bovine serum (Peak Serum, #PS-FB2, Lot #21E1202), 2 mM glutamine (Corning, #25-005-CI), and penicillin/streptomycin (Corning, #30-002-CI).

SYTOX Green (#S7020) and LIVE/DEAD Fixable Violet Dead Cell Stain (#L34964) were purchased from Thermo Fisher Scientific. p53 mouse antibody (#48818) and p-H2A.X rabbit antibody (#9718) were purchased from Cell Signaling Technologies. β-Actin mouse antibody (#A2228) and actin rabbit antibody (#A2066) were purchased from Sigma-Aldrich. Anti-Active Caspase-3 antibody (#559565) and Anti-Cleaved PARP-647 (#558710) were obtained from BD Pharmingen. Goat anti-rabbit Alexa-488 secondary antibody (#A-11008) was purchased from Thermo Fisher Scientific. Total OXPHOS Rodent WB Antibody Cocktail was purchased from Abcam (ab110413). Anti-phospho-Histone H3 rabbit antibody (#06-570) and propidium iodide (#81845) were purchased from MilliporeSigma. Camptothecin (#S1288), Carboplatin (#S1215), Chlorambucil (#S4288), Cisplatin (#S1166), Idarubicin HCl (#S1228), Irinotecan (#S2217), Etoposide (#S1225), Ferrostatin-1 (#S7243), Necrostatin-1 (#S8037), Nutlin-3 (#S1061), Rucaparib (#S1098), Teniposide (#S1787), and Topotecan HCl (#B2296) were purchased from Selleck Chemicals. ABT-199 (#A8194), E 64D (#A1903), Hydroxychloroquine Sulfate (#B4874), VX-765 (#A8238), Z-IETD-FMK (#B3232), and Z-VAD-FMK (#A1902) were purchased from APExBIO. Cyclosporin A (#30024) was obtained from Sigma-Aldrich. Blasticidin (#BP264750) was bought from Fisher Scientific, and Puromycin (#61-385-RA) was purchased from Corning. Pooled siGENOME siRNAs were purchased from Dharmacon/Horizon Discovery. The non-targeting siRNA pool (#D-001206-13-05) contains sequences UAGCGACUAAACACAUCAA, UAAGGCUAUGAAGAGAUAC, AUGUAUUGGCCUGUAUUAG, and AUGAACGUGAAUUGCUCAA. Pooled siRNAs against p53 (#M-003329-03-0005) include the sequences GAGGUUGGCUCUGACUGUA, GCACAGAGGAAGAGAAUCU, GAAGAAACCACUGGAUGGA, and GCUUCGAGAUGUUCCGAGA.

### Immunoblotting

Lysates for immunoblotting were prepared from cells seeded in either 6-well plates (200,000 cells per well) or 10 cm dishes (1.5×10^6^ cells per dish). Cells were adhered overnight and then drugged the following morning (“T0”). At each of the indicated timepoints, media was removed from each sample and collected in a conical tube. Samples were washed with PBS, which was collected in the same conical tube. Cells were then trypsinized and pelleted together with their corresponding media/PBS wash. SDS-lysis buffer (50 mM Tris-HCl, 2% SDS, 5% glycerol, 5 mM EDTA, 1 mM NaF, 10 mM β-glycerophosphate, 1 mM PMSF, 1 mM Na_3_VO_4_, protease inhibitor tablet and phosphatase inhibitor tablet) was used to lyse the cell pellets. Centrifugation through a 0.2 μm multi-well filter plate (Pall Laboratory, #5053) was used to remove DNA from the lysates. The protein concentration of each sample was quantified with a BCA assay (Thermo Fisher Scientific, #23225). Lysates were normalized to the same protein concentration and run on precast 48-well E-PAGE gels (Thermo Fisher Scientific, #EP04808). Gels were transferred to nitrocellulose membranes using the iBlot semidry system (Invitrogen) and then blocked for 1 hour in 50% PBS: 50% Odyssey Blocking Buffer (OBB, LI-COR Odyssey, #927-40010). Membranes were incubated overnight at 4°C in primary antibody (diluted in 50% PBS-T (PBS + 0.1% Tween-20): 50% OBB), and then stained with infrared dye-conjugated secondary antibodies (LI-COR). A LI-COR Odyssey CLx scanner was used to visualize the immunoblots.

### Evaluation of p53 function

For our re-analysis of public data, we divided cell lines into p53-proficient and p53-deficient groups based on their DepMap annotated sensitivities to MDM2 deletion. MDM2 inhibition causes a proliferation defect only for cells that retain p53 function^65^. For this reason, MDM2 deletion scores as “essential” in screens of p53 WT cells. Based on this analysis, we identified 90 cell lines that have functional p53, and 305 cell lines in which p53 is non-functional. This classification was used to determine the DNA damage sensitivity of p53 WT and p53 KO cell lines from DepMap. The drug sensitivity data used was the PRISM Repurposing Primary Screen, version 19Q4. This data comes from a multiplexed cell-line viability assay that was used to evaluate a large panel of small molecules. Each cell line in the dataset was bifurcated by p53 status, and drugs whose mechanism of action is DNA damage were identified. The distribution of PRISM fold-change scores for these drugs was plotted as a violin in Prism 9.

Functionality of p53 in our isogenic cell lines was assessed with two complementary methods. First, expression of p53 was measured using a western blot. Stabilization of p53 was induced by treating cells with 10 µM Camptothecin for 2 hours, and immunoblotting was performed with an antibody against p53. Second, flow cytometry was used to determine the functionality of p53. U2OS cells and U2OS^p53KO^ clones were treated with 10 µM Nutlin-3 for 24 hours, and stained with PI and pH-H3 antibody (described in detail below). Measurement of cell cycle position and cell cycle arrest was performed in FlowJo, and cells without p53-dependent G1 and G2 checkpoints were classified as p53-deficient (KO).

### Generation of U2OS*^p53KO^*

U2OS*^p53KO^* cells were generated in two steps. First, U2OS cells were transduced with virus containing lentiCas9-Blast (Addgene, #52962). Cells were then treated with 5 µg/mL blasticidin for 5 days to make a stable population of U2OS-Cas9 cells. A gRNA cloning vector (Addgene, #41824) was then used to knock out p53. The synthesis protocol provided by the Church lab (available on Addgene) was followed to generate a plasmid with an sgRNA against p53 (5’GATCCACTCACAGTTTCCAT’3). The p53-sgRNA plasmid was transiently transfected into U2OS-Cas9 cells using Lipofectamine 2000 (Thermo Fisher Scientific, #11668027). The transfected cells were then treated with 10 µM Nutlin-3 for 7 days to enrich for p53 KO cells. Nutlin-3 disrupts the interaction of p53 and MDM2, forcing cells with wildtype p53 to engage cell cycle arrest. p53 KO cells continue to grow in the presence of Nutlin-3 and become enriched in the population over time. From this highly enriched population of p53 KO cells, single cells were cloned and then tested for loss of p53 using western blotting and Sanger sequencing (sequencing primers 5’GCTGGATCCCCACTTTTCCTCT’3 and 5’CATCCCCAGGAGAGATGCTGAG’3).

### Assays to measure drug-induced cell death

The FLICK assay was performed as described in Richards et al. 2020^9,26^. Briefly, cells were seeded at a density of 2,000-5,000 cells per well in black 96-well plates (Greiner, #655090). Cells were plated in 90 µL of media and incubated overnight. Cells were then drugged with the indicated doses of drug or vehicle controls, along with 5 µM SYTOX green, in 10 µL media. Dead cell fluorescence was monitored kinetically with a Tecan Spark microplate reader (ex:503, em:524), using a gain that achieved linearity of the SYTOX signal for each cell line. A duplicate plate was lysed at the beginning of the assay by adding 0.1% Triton X-100 (Fisher Scientific, #BP151-100) in PBS to each well, and incubating each plate at 37°C for 2-4 hours. At the end of the experiment, all wells were permeabilized with 0.1% Triton X-100 in PBS. The permeabilized plates were used to determine the total cell number at the assay start and endpoint. Total cell number at intermediate timepoints was inferred using a simple model of exponential growth. The measured dead cell fluorescence and total cell fluorescence was then used to calculate the fluorescence of live cells at each timepoint. From these 3 numbers, all calculations necessary to determine relative viability (RV), fractional viability (FV), and GR value (GR) can be made.

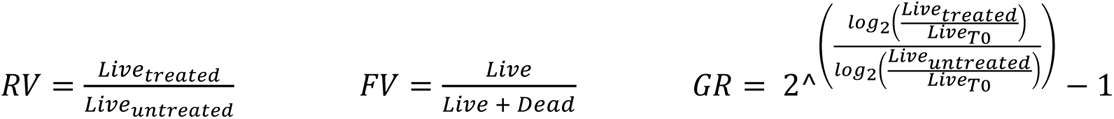

Dose response curves and LF kinetic curves were calculated using a custom MATLAB script, as described previously (Richards et al., 2020). Sigmoidal dose response curves were fit using a four-parameter logistic regression to determine the plateau, hill slope, EC_max_, and EC_50_. These parameters were determined using least squares regression-based curve fitting in MATLAB. An example is shown below for FV, but the same equation was also used to fit RV and GR.

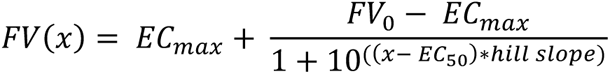

The previously described lag-exponential death (LED) equation was used to model cell death over time^33^. This model optimizes several parameters, including LF_0_ (the starting lethal fraction), LF_p_ (the death plateau), D_0_ (the death onset time), and D_R_ (the maximum rate of death). The area under the resulting LED curve was also calculated (AUC).

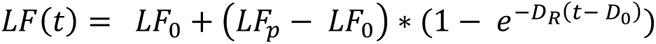

Drug GRADE was calculated as described in Schwartz et al., 2020^27^. FV and GR were calculated as described above, and then FV values were normalized relative to the basal death rate of each cell line. For each drug, the GR and FV from doses where GR ≥ 0 were fit to a line to determine the slope (*m_drug_*). The maximum slope that could be observed over the same range of GR values was also determined (*m_max_*), and then GRADE was calculated.

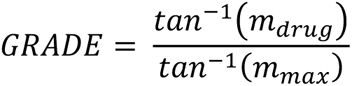

### Flow cytometry-based analysis of apoptosis and cell cycle

Cleaved-CASP3 and cleaved-PARP positivity was quantified to monitor activation of apoptosis. At the indicated timepoints, the media was collected from each sample and the remaining adherent cells were trypsinized. The media and trypsinized cells were pooled for each sample, pelleted, and then stained with a 1:1000 dilution of LIVE/DEAD fixable violet stain at room temperature for 30 minutes. Each sample was then washed with cold PBS. Samples were fixed in 4% fresh formaldehyde in PBS at room temperature for 15 minutes. Cells were washed with cold PBS, pelleted, and resuspended in ice-cold 100% methanol. The fixed and permeabilized cells were then stored at –20°C overnight. The methanol was removed the following day, and samples were washed twice with PBS-T (PBS + 0.1% Tween-20). Primary cleaved-CASP3 antibody was diluted 1:500 in 50% PBS-T: 50% Odyssey Blocking Buffer (OBB, LI-COR Odyssey, #927-40010). Samples were then incubated with diluted cleaved-CASP3 antibody at room temperature for 8 hours. After incubation, samples were washed once with PBS-T, and then incubated with cleaved-PARP-647 antibody and goat-anti rabbit Alexa-488. Both antibodies were diluted 1:250 in 50% PBS-T: 50% OBB and incubated overnight at room temperature. Samples were then pelleted, washed twice with PBS-T, resuspended in PBS-T, and filtered in preparation for flow cytometry analysis. Samples were run on either a BD LSRII or a Miltenyi MACSQuant VYB cytometer with laser and filter settings appropriate for reading LIVE/DEAD violet, Alexa-488, and Alexa-647. Analysis to identify live cells and quantify the number of cleaved-CASP3+/cleaved-PARP+ cells was performed using FlowJo.

For the analysis of cell cycle position, media and adherent cells were collected as described above. Cells were pelleted and then fixed by gentle resuspension in ice-cold 70% ethanol in PBS. Cells were then stored at –20°C overnight. Each sample was then washed twice with PBS, and permeabilized on ice for 15 minutes using Triton x-100 (0.25% in PBS). Permeabilized cells were then rinsed with 1% BSA in PBS. Samples were then incubated overnight at 4°C with phospho-histone H3 antibody, which was diluted 1:100 in 1% BSA. The following day, cells were washed twice with 1% BSA and then incubated with goat-anti rabbit Alexa-488 secondary (in 1% BSA) for 1 hour at room temperature. Cells were then washed once with 1% BSA and once with PBS. Each sample was then resuspended in 10% RNase A in PBS (Sigma-Aldrich, #R6513). Propidium iodide (10mg/ml) was then added to each sample at a final concentration of 0.5mg/ml. Samples were then filtered and analyzed on a BD LSRII. Cell cycle position was quantified using FlowJo analysis software.

### Imaging and time-lapsed microscopy

For imaging experiments to monitor apoptotic/non-apoptotic morphologies, U2OS and U2OS*^p53KO^* cells were plated in black 96-well plates. Cells were plated at a density of 2,000-5,000 cells per well, depending on the goals of the experiment. Cells were allowed to adhere overnight and then treated with the indicated drugs and 50 nM SYTOX green. Endpoint or time-lapsed images were then collected using either an IncuCyte S3 or an IncuCyte SX5 (Essen Biosciences). Images were collected at 20× magnification in phase, as well as the green channel (ex: 460 ± 20, em: 524 ± 20, acquisition time: 300ms). Images were visualized and analyzed using Fiji (ImageJ2).

Time-lapsed microscopy was also utilized in the STACK assay. Prior to the assay, U2OS-mCherry and U2OS^p53KO^-mCherry cells were generated. Integration of mCherry into the genome of these cells was achieved using a viral H2B-mCherry plasmid. Cells were then plated and drugged as above. Images were collected on the IncuCyte S3 at 10× magnification in phase, the green channel (ex: 460 ± 20, em: 524 ± 20, acquisition time: 300ms), and the red channel (ex:585 ± 20, em: 635 ± 70, acquisition time: 400ms). The counts per well for the SYTOX+ and mCherry+ objects were then determined using the built-in IncuCyte Software (Essen Biosciences) and exported to excel for analysis using a custom MATLAB script.

### RNA-seq of conditioned media and evaluation by GSEA

Conditioned media was generated from U2OS and U2OS^p53KO^ cells, before and after exposure to etoposide. For both genotypes, 1×10^6^ cells were plated on 10 cm plates and incubated overnight. Cells were then treated with either 31.6 µM etoposide or DMSO. After 48 hours of drug or vehicle treatment, the conditioned media was collected and passed through a 0.45 µm filter to remove cellular debris. Each conditioned media was diluted 50:50 with fresh media to replenish growth factors, and then immediately used to treat U2OS cells. The recipient U2OS cells were plated at 300,000 cells/well in a 6-well plate 1 day prior to treatment. A T0 sample was also collected prior to addition of the conditioned media. U2OS cells were exposed to the conditioned media for 8 hours and then trypsinized and flash frozen. Total RNA was extracted using the RNeasy Mini Kit (Qiagen, #74104). The manufacturer’s instructions were followed to purify RNA, and then 25×10^6^ million reads were sequenced for each sample. RNA sequences were aligned and counted using the DolphinNext RNA-seq Pipeline^66^. Genes with less than 20 counts were trimmed, and DESeq2’s parametric fit was used to calculate the log_2_ fold-change (L2FC) and adjusted p-value for each gene. Hits were identified based on both a L2FC (x<-0.4 or x>0.4) and significance cutoff (x<0.05). GSEA was run with a pre-ranked list of the L2FC values to determine pathway-level enrichments. Raw data files associated with this experiment are available from the Gene Expression Ombibus (GEO, accession numbers: GSM7356394-7356398; GSM7356400-7356404).

### BH3 profiling to evaluate apoptotic priming

U2OS or U2OS*^p53KO^* cells were collected through trypsinization and pelleting, and then subjected to BH3 mimetic profiling as described previously^67,68^. In brief, cells were permeabilized with digitonin and then treated with activator or sensitizer BH3 peptides in Mannitol Experimental Buffer (10 mM Hepes (pH 7.5), 150 mM mannitol, 50 mM KCl, 0.02 mM EGTA, 0.02 mM EDTA, 0.1% BSA, and 5 mM succinate). Alamethicin was used as a positive control, and a mutant inactive PUMA peptide (PUMA2A) was used as a negative control. After 60 minutes of peptide exposure, cells were fixed and stained overnight with an antibody for cytochrome c in a saponin-based intracellular staining buffer. Cytochrome c release was then measured using an Attune NxT flow cytometer.

### Simulation of population-level responses to DNA damage in the presence and absence of p53

The response of cells to DNA damage was modeled as a biphasic response. For untreated cells, growth is modeled simply as an exponential growth equation (equation I). For cells treated with DNA damage, the change in population size is modeled with exponential equations for growth and death (equation II).

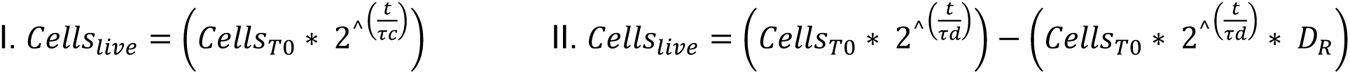

To simulate a dose curve of treated p53 WT cells, the modeled growth rate was set to 0, and the drug-induced death rate was parameterized around experimentally observed values. To simulate how loss of cell cycle arrest would affect the population size, the growth rate of treated cells was parameterized using the experimentally observed growth rate in p53 KO cells treated with DNA damage (Figure S2B), while maintaining the experimentally observed p53 WT death rates. Loss of cell death was simulated by setting the death rate to 0. Code used to simulate RV in the presence/absence of p53-dependent growth inhibition and/or death activation is available on GitHub (https://github.com/MJLee-Lab/RVsim).

### Chemo-genetic profiling of responses to DNA damage in the presence and absence of **p53**

Whole-genome CRISPR screens were performed in U2OS and U2OS*^p53KO^* cells using the GeCKOv2 two-vector system (Addgene, #1000000049). The two pooled DNA half-libraries (A and B) were combined to generate a library of 123,411 sgRNAs. This library was amplified according to the Zhang lab’s protocol (available on Addgene) and virus was generated using 293T cells. U2OS-Cas9 and U2OS*^p53KO^*-Cas9 cells were generated using viral transduction of the lentiCas9-Blast plasmid (Addgene, #52962), followed by a 5-day selection with 5 µg/mL blasticidin. U2OS-Cas9 and U2OS*^p53KO^*-Cas9 cells were transduced with the GeCKOv2 viral library using a “spinfection”. For each genotype, >200 x10^6^ cells were spinfected to achieve a total library coverage of 300-500×. To perform the spinfection, 2×10^6^ cells were combined with 300 µL of virus and 0.8 µL/mL polybrene (Millipore, #TR1003G) in the well of a 12-well plate (final volume of 2 mL). This volume of virus was selected experimentally to achieve a final MOI of 0.3. The 12-well plate was then centrifuged at 37°C for 2 hours at 830 x g. The media was gently replaced after the spin and cells were allowed to recover overnight. The following day, cells were replated into 8-layer flasks (Greiner Bio-one, #678108). Cells were incubated overnight and then treated with 1 µg/mL puromycin for 3 days. After the 3-day puromycin treatment the cells were replated into 8-layer flasks and further expanded for 3 days. On day –1 of the drug screen, 50-100 x10^6^ cells (∼400 – 800× coverage) were plated in triplicate for each of the experimental conditions. For each genotype, 50 x10^6^ cells were also saved in triplicate for the T0 controls. On day 0, the treated conditions were drugged with 5 µM etoposide. Untreated cells were passaged on days 1 and 3 (maintaining 400× coverage), and treated cells were passaged on day 3. Live cells from the treated and untreated conditions were all collected and frozen on day 4. Genomic DNA was isolated from the cell pellets using a phenol-chloroform– based extraction method, and sgRNA sequences were extracted from each genome by PCR (forward: 5’CGATTTCTTGGCTTTATATATCTTGTGG’3 and reverse: 5’CTCTGCTGTCCCTGTAATAAACC’3). A second round of PCR was used to add multiplexing barcodes, and each gel-purified library was sequenced on a HiSeq4000 at 300× coverage. Raw data files are available from on the Gene Expression Omnibus (GEO; accession numbers 7356374, 7356376-7356385, 7356387-7356393).

### Analysis of chemo-genetic profiles using relative population size (L2FC)

The read quality of the sequences from the chemo-genetic screen were verified using FastQC, and then the libraries were de-multiplexed using the barcode splitter function in the FASTX tool kit. The FASTX trimmer function was used to remove the non-variable sgRNA regions from each sequence. Reads were mapped to the GeCKOv2 library using Bowtie2, allowing for a single mismatch. Guides with low counts were trimmed by removing the bottom 5% of sgRNAs. miRNA sequences were also removed. Sequencing depth was normalized using the distribution of the 1000 non-targeting guides. For each comparison of interest (untreated/T0 and treated/untreated for each genotype), the log_2_ fold-change (L2FC) was calculated using a parametric fit in DESeq2. The GeCKOv2 library contains 6 sgRNAs for each of 19,050 genes. These 6 guide-level scores were collapsed to a single gene-level L2FC by taking the mean. The non-targeting sgRNAs were randomized and assigned to 6-guide non-targeting “genes”. Each fold-change value was then z-scored based on the distribution of L2FC scores for the non-targeting genes. An empiric p-value was determined for each gene, and this score was FDR corrected using the Benjamini-Hochberg procedure.

### Death rate-based analysis of chemo-genetic profiles

Conventional methods for analyzing whole-genome CRISPR screens, which compare the relative population size of treated and untreated conditions (L2FC), are dominated by “essential” pathways (Figure 3E-G). Like relative viability, L2FC is highly sensitive to changes in growth rate and insensitive to changes in the death rate. Based on this observation, it was clear that accurately identifying death regulatory genes from our chemo-genetic profiling data would require an analysis strategy that scores changes in the drug-induced death rate. We began by developing a simple model of population dynamics in the presence and absence of DNA damage (Figure 4A). Growth and death rates for our model were parameterized based on drug titration data used to optimize our screen (Figure 3B-C). Using this model, we simulated all possible combinations of growth rates and drug-induced death rates, treating these features as independent variables. From the simulated data, we computed the relative size of the treated and untreated populations, as is conventionally done in fold change-based analysis of chemo-genetic profiles. The results of this comprehensive simulation revealed with more clarity the nature of the confounding influence of growth rate variation: it is possible to accurately interpret the direction and magnitude of each gene deletion’s effect on cell death, but only if the gene deletion does not result in a growth rate perturbation (Figure 4B). For each single gene knockout, as the growth rate in the untreated condition is decreased from the wild-type growth rate, the inference of the gene’s function in the treated condition is compromised, even inverted if the growth defect is strong enough (Figure 4C). For example, knocking out DNA repair genes significantly compromises the growth rate of cells, even in the absence of external DNA damage (Figure 4D). This slow growth in the untreated condition is responsible for the inverted inference of DNA repair genes (Figure 4E-F).

Our simulation also revealed a straight-forward strategy for inferring the drug-induced death rates from our existing data. The central issue is that the population size at assay endpoint is not proportional to the drug-induced death rate, but rather, is a function of the genetic-and drug-induced changes to both growth and death rates. Under the conditions tested in our chemo-genetic screen, DNA damage-induced death occurs only after cells have stopped proliferating (Figure S5A). The lack of growth during the death phase of the response significantly simplifies the possible ways that perturbations to growth rate and death rate integrate to create a given number of cells. Thus, the varied combinations of growth rates and death rates that yield the same population size create a single continuous non-linear “manifold” at each level of L2FC (Figure S5B). The implication of this simple structure is that the drug-induced death rate for each single gene knockout can be clearly inferred from a combination of the relative population size (e.g., L2FC) and the relative growth rate in the absence of drug (e.g., the gene fitness score when comparing untreated to T0) (Figure S5C).

To determine the drug-induced death rate for each sgRNA, we first parameterized the model described above around the observed growth and death rates of wild-type U2OS cells (Figure 3C-D). The second measurement that is required for this analysis is the experimentally observed L2FC for each sgRNA. The L2FC for etoposide/untreated and untreated/T0 was computed as described above. To identify which combination of growth rate and death rate generated the observed L2FC, the growth rate of each SGKO was experimentally determined by comparing the untreated cells to the T0 input sample. From the relative population size (*i.e.*, L2FC) in etoposide/untreated conditions, combined with the inferred growth rate of each clone, and the observed coordination of growth and death in GRADE-based analysis, a high-confidence inference of the drug-induced death rate for each knockout clone can then be generated (Figure S5C). This procedure was used to determine the relative growth rate and relative drug-induced death rate of each sgRNA. Similar to the procedure for calculating L2FC, the 6 guide-level scores for each gene were collapsed by taking the mean. The gene-level data for growth and death rate can then be projected into the phase diagram generated from our simulation (Figure 4E). To generate a p-value, the gene-level growth and death rates were z-scored based on the distribution of rates for the non-targeting genes. The z-scored values were bootstrapped to determine an empiric p-value and FDR corrected. Full code used to estimate growth and death rates for single gene knockouts in the context of DNA damage is available on GitHub (https://github.com/MJLee-Lab/DRADD).

### Screen validation

The chemo-genetic CRISPR screen and our rate-based analysis were validated in p53 wildtype cells treated with etoposide. This validation effort included the top and bottom 10 genes from the fold-change based analysis, the top and bottom 10 genes from the death rate-based analysis, and 4 genes involved in the canonical repair of etoposide-induced DNA damage. For each gene, the highest-scoring sgRNA was selected from the GeCKOv2 library. Each guide was cloned into the pX330-puro plasmid. Cloning was performed using the single-step digestion-ligation protocol from the Zhang lab (available on the Zhang lab Addgene page). To validate each guide, U2OS cells were plated in 6-well plates at a density of 200,000 cells per well. Lipofectamine 3000 (Thermo Fisher Scientific, #L3000008) was used to transiently transfect each sgRNA-pX330-puro plasmid into a separate well. Cells were treated with 1 µg/mL puromycin for 3 days after the transfection. Cells were replated at the end of the antibiotic selection and allowed to recover for 2 days. Post-recovery, each pool of single gene knockouts (SGKOs) was plated at 2,000 cells per well on black 96-well plates. Changes in drug sensitivity and growth rate were then measured using FLICK. For measurement of death rate, cells were treated with 5 µM etoposide for 4 days. The drug sensitivity of each SGKO pool was determined by comparing the LF of each sgRNA to that of a non-targeting sgRNA control (ΔLF) at 96 hours. Non-targeting values across 4 separate experiments were averaged for robustness. Fisher’s exact tests were used to evaluate the performance of the fold-change and rate-based analysis methods. For measurement of growth rate, plates of untreated cells were lysed at 24, 48, and 72 hours with 0.1% Triton X-100. The total cell number at each timepoint was fit to an exponential model of growth to determine the growth rate of the non-targeting controls and the SGKO pools.

### Cobalt-calcein assay for mPTP

The stains used in the mPTP assay (calcein, cobalt (II) chloride hexahydrate, and MitoTracker Red) were obtained from the Image-iT LIVE Mitochondrial Transition Pore Assay Kit (Thermo Fisher Scientific, #I35103). In preparation for measurement of MPT activation, cells were seeded on round glass coverslips in 12-well plates. Treated wells were seeded at a density of 100,000 cells/well, and untreated cells were seeded at 30,000 cells/well. After overnight adherence, cells were treated with the indicated drugs. Cells were stained with calcein at a timepoint after the onset of cell death (as measured by FLICK). Coverslip-adhered cells were washed with PBS supplemented with 1 mM CaCl_2_ (PBS-Ca). PBS-Ca containing 0.5 µM calcein and 0.2 µM MitoTracker Red was then added to each sample. Samples were then spiked with 10 mM CoCl_2_ and mixed by pipetting. Cells were then incubated at 37°C in 5% CO_2_ for 15 minutes. Samples were washed twice with PBS-Ca and then submerged in PBS-Ca until imaging. Each glass coverslip was placed on a slide immediately prior to imaging, and all images were taken within 30 minutes of staining. Images were collected on an EVOS FL Auto 2 microscope with a 40× objective using GFP (ex:470, em:510) and Texas Red (ex:585, em:624) light cubes (Life Technologies). Images were analyzed in Fiji (ImageJ2). Each cell was masked based on its MitoTracker Red signal, and then the mean fluorescence intensity of calcein was determined for each cell. A minimum of 50 cells were counted for each experimental condition.

### Seahorse Extracellular Flux Assay

Real-time oxygen consumption rate (OCR) was measured using a Seahorse XF96 Extracellular Flux Analyzer (Agilent) and a Mito Stress Test Kit (Agilent, #103015-100) according to the manufacturer’s instructions. U2OS or U2OS*^p53KO^* cells were seeded in XF96-well cell culture microplates (Agilent, #101085-004) at 30,000 cells/well in high glucose and incubated at 37°C in 5% CO_2_ overnight. For the indicated timepoints, cells were drugged with 31.6 µM etoposide. Timepoints were drugged in reverse (i.e., 16, 8, 4, 2, 0 hours) so that the OCR could be read for all conditions simultaneously. Prior to performing the Mito Stress Test, growth medium was replaced with 180 µL of DMEM (without serum) supplemented with 25 mM glucose and 2 mM L-glutamine. Cells were incubated at 37°C for 60 minutes, and then four baseline OCR measurements were taken before adding the assay compounds. Mitochondrial respiration was assessed using 1 µM oligomycin, 1 µM FCCP, and 1 µM rotenone/1 µM antimycin. For each phase of the Mito Stress Test 3 measurements were taken. Following measurements of OCR, the total cell number for each well was determined using a modification of the FLICK assay. SYTOX Green was added to each well of the Seahorse plate at a final concentration of 15 µM, and cells were lysed for 4 hours using 0.15% Triton X-100. In parallel, a titration of U2OS cells was plated in a black 96-well plate and lysed with Triton X-100. The Seahorse plate and cell titration plate were read on a Tecan Spark at a gain of 100. The fluorescence of 30,000 U2OS cells was determined from the cell titration, and this value was used as a normalization factor for the measured fluorescence of the Seahorse plate. Respiration parameters (non-mitochondrial oxygen consumption, basal respiration, maximal respiration, proton leak, ATP production, and spare respiratory capacity) were then calculated for each condition across 3-6 replicates.

### Electron microscopy

U2OS and U2OS*^p53KO^* cells were grown in fully supplemented growth medium on 10 cm dishes prior to collection for transmission electron microscopy (TEM). Adhered cells were fixed using a 50:50 mixture of growth medium and fixation solution (2.5% glutaraldehyde and 1.6% paraformaldehyde in 0.1M sodium cacodylate buffer, pH 7.4) for >30 minutes, and then moved to 100% fixation solution for 1 hour. After fixation, cells were washed with 0.1M sodium cacodylate buffer and then post-fixed in osmium tetroxide. Cells were scraped and pelleted, and then dehydrate through a graded ethanol series. Cell pellets were treated with propylene oxide and infiltrated in SPI-pon/Araldite for embedding. Ultra-thin sections were cut on a Leica UC7 Ultramicrotome and imaging was performed using a Phillips CM120 Transmission Electron Microscope.

### Evaluation of mitochondrial abundance

#### MT-DNA

Mitochondrial DNA (MT-DNA) was quantified using qPCR. Genomic DNA and MT-DNA was extracted using a DNeasy Blood & Tissue Kit (Qiagen, #69504) according to the manufacturer’s instructions. Total DNA mass was quantified using a Qubit 4 (Thermo Fisher Scientific) and a dsDNA BR Assay Kit (Thermo Fisher Scientific, #Q32850). MT-DNA was then quantified on a QuantStudio 3 Real-Time PCR System (Thermo Fisher Scientific) using Power SYBR Green PCR Master Mix (Thermo Fisher Scientific, #4367659) and the following primers: MT-DNA marker D-loop (Forward: 5’TATCTTTTGGCGGTATGCACTTTTAACAG’3, Reverse: 5’TGATGAGATTAGTAGTATGGGAGTGG’3), and nuclear DNA marker B-Actin (Forward: 5’TCACCCACACTGTGCCCATCTACGA’3, Reverse: 5’CAGCGGAACCGCTCATTGCCAATGG’3). D-loop primers were used at a final concentration of 1 µM, B-Actin primers were used at a concentration of 250 nM, and 0.2 ng of total DNA was used for each reaction. Both primer pairs and DNA concentration were optimized to achieve high-efficiency exponential amplification of the targets. Thermo-cycling conditions were as follows: initial denaturation (1 min at 95°C), cycling stages (15s at 95°C, 30s at 61°C, 30s at 72°C, x 40 cycles), and melt curve (1 min at 60°C and then a 0.3°C/s ramp down from 95°C). Delta CT was determined for each genotype using B-Actin as the control gene in Thermo Fisher Scientific’s Design & Analysis Software, Version 2.6.0.

#### Mitochondrial mass

Mitochondrial mass was quantified using MitoTracker Green FM (Thermo Fisher Scientific, #M7514). U2OS and U2OS*^p53KO^* cells were pelleted and washed with PBS, and then resuspended in 150 nM MitoTracker in PBS + 5% FBS. Cells were incubated for 25 minutes at 37°C, washed, and immediately analyzed on a Miltenyi MACSQuant VYB cytometer with laser and filter settings appropriate for reading FITC. Analysis was performed using FlowJo.

### Electron transport chain complex abundance and SC formation

#### Immunoblotting

Lysates for denatured gels were prepared from cells seeded in 10 cm dishes. Untreated U2OS and U2OS*^p53KO^* cells were collected as described above through trypsinization. Cells were lysed using a standard RIPA buffer (150 mM NaCl, 50 mM Tris pH 7.5, 15 mM Na_4_P_2_O_7_, 50 mM NaF, 1 mM EDTA, 1 mM Triton X-100, 10 mM β-glycerophosphate, 2 mM sodium orthovanadate, 1 mM DTT, protease inhibitor tablet, and phosphatase inhibitor tablet). After pelleting cellular debris, supernatant was quantified using a BCA assay (Thermo Fisher Scientific, #23225). Protein concentration was normalized and run on 10% polyacrylamide gels, and then transferred to a nitrocellulose membrane. Membranes were blocked for 1 hour in 50% PBS: 50% Odyssey Blocking Buffer (OBB, LI-COR Odyssey, #927-40010). Membranes were then incubated overnight at 4°C in primary antibody (diluted in 50% PBS-T (PBS + 0.1% Tween-20): 50% OBB). The Total OXPHOS Rodent WB Antibody Cocktail from abcam (ab110413) was used at a dilution of 1:1000 to simultaneously measure all 5 ETC complexes. Following overnight incubation in primary antibody, membranes were stained with infrared dye-conjugated secondary antibodies (LI-COR). A LI-COR Odyssey CLx scanner was used to visualize the immunoblots.

#### Isolation of mitochondria for native gels

Mitochondria were isolated from 3 separate biological replicates of U2OS and U2OS*^p53KO^*. Each replicate contained 50-70 x 10^6^ cells, harvested from multiple 15 cm plates. Cells were pelleted, washed with PBS, and resuspended in 0.5 mL of ice-cold isolation buffer (200 mM sucrose, 10 mM Tris, 1 mM EGTA/Tris, adjusted to pH 7.4 with 1M HEPES). Cell suspensions were syringe lysed with 6 fast strokes.

Homogenate was then spun at 600 x g for 10 minutes at 4°C to separate cell debris. The supernatant from this spin was then transferred to a new tube and centrifuged at 7,000 x g for 10 minutes at 4°C. The supernatant was discarded, pelleted mitochondria were gently washed with 1 mL of cold isolation buffer, and mitochondria were pelleted again at 7,000 x g for 10 minutes at 4°C. Mitochondrial protein was then quantified using a BCA, and pellets were stored at –80°C until ready for use.

#### Blue native PAGE

Blue native PAGE (BN-PAGE) gels were run as previously described^69^, using mitochondria isolated as described above. Mitochondrial protein (50 µg) was re-suspended in 1X NativePAGE Sample Buffer (Thermo Fisher Scientific, #BN2003) and 4 g digitonin:1 g mitochondrial protein (Sigma-Aldrich, #D141-100MG). Mitochondria were incubated on ice for 20 minutes and then centrifuged at 20,000 x g for 10 minutes at 4°C. Coomassie NativePAGE 5% G-250 Sample Additive (Thermo Fisher Scientific, #BN2004) was combined to 1/4^th^ the final protein concentration in the supernatant. Samples were then loaded onto NativePAGE 3 to 12% Bis-Tris Mini Gels (Thermo Fisher Scientific, #BN1003BOX) along with NativeMark Unstained Protein Standard (Thermo Fisher Scientific, #LC0725). The inner chamber of the gel apparatus was filled with dark blue cathode buffer, and the outer chamber was filled with anode buffer from the NativePAGE Running Buffer Kit (Thermo Fisher Scientific, #BN2007). Gels were run for 30 minutes at 150 volts, and then the dark blue cathode buffer was replaced with light blue cathode buffer from the NativePAGE Running Buffer Kit. Gels were then run for an additional 150-170 minutes at 250 volts to separate ETC complexes. Each gel was then transferred to a 0.45 µm PVDF membrane using 10% methanol-supplemented 1X NuPAGE Transfer Buffer (Thermo Fisher Scientific, #NP0006) for 1 hour at 30 volts. Membranes were washed with 8% acetic acid, dried, and de-stained with 100% methanol. Blocking was performed with 5% milk in 1X TBST for 1 hour, and primary antibody incubation was performed in 5% BSA with rocking overnight at 4°C. Total OXPHOS Antibody Cocktail was used as described above. Blots were washed with TBST, and membranes were incubated with (HRP)-linked secondary antibody in 5% milk for 1 hour at room temperature. Membranes were washed and then developed using ECL substrate (Thermo Fisher Scientific, #32106).

### Metabolite profiling

#### Metabolite isolation and LC-MS

Cells were seeded on 6-well plates at 200,000 cells per well. After adherence overnight, cells were treated with DMSO, 5 µM etoposide, or 31.6 µM etoposide. At the indicated timepoints, samples were collected in parallel for metabolite extraction or protein quantification. For total protein quantification, cells were trypsinized and pelleted. Pellets were lysed using SDS lysis buffer and quantified using a BCA assay (described above). For extraction of metabolites, growth media was removed, and cells were washed 2 times with ice-cold PBS. With the 6-well plate on dry ice, cells were submerged in 500 µL 80% MeOH. Samples were then incubated at –80°C for 15 minutes. Cell scrapers were used to harvest each sample, and sample wells were washed with an additional 300 µL of 80% MeOH. Samples were vortexed at 4°C for 10 minutes and then centrifuged at top speed for 10 minutes at 4°C. Supernatant was transferred to a new tube and samples were dried using a speed vac. Dried pellets were resuspended in 100 µL of water and vortexed for 10 minutes at 4°C. Samples were then spun for 10 minutes at top speed at 4°C, and supernatant from each sample was transferred to an LC-MS vial (Thermo Fisher Scientific, #6ESV9-04PP and #6ASC9ST1). A QExactive Plus Quadrupole Orbitrap Mass Spectrometer equipped with a HESI II probe (Thermo Fisher Scientific) was then used to perform Mass Spectrometry. Metabolites were quantified by integrating peaks in TraceFinder 5.1 (Thermo Fisher Scientific). Mass tolerance was set to 5 ppm and expected retention times were benchmarked using an in-house library of chemical standards. Raw ion counts were normalized to the total protein amount, as quantified using BCA. Fold-change and significance was determined using a custom MATLAB script. Enriched metabolic signatures were identified using Pathway Analysis in MetaboAnalyst 5.0. Raw data are available for download at DOI 10.5281/zenodo.7931639.

#### Isotope tracing using ^13^C_6_-Glucose

Isotope tracing experiments were performed using heavy D-^13^C_6_-Glucose (Cambridge Isotope Laboratories, #CLM-1396). Cells were seeded in 6-well plates, incubated overnight, and treated with drug as above. Eight hours prior to the collection of each timepoint, the growth medium was swapped with glucose-free DMEM (Thermo Fisher Scientific, #11966025) supplemented with 10 mM D-^13^C_6_-Glucose. Metabolites were extracted from cells using 80% methanol as above, and metabolite abundance was quantified using LC-MS and TraceFinder. Natural abundance of heavy isotopes was corrected using IsoCorrectoR (Bioconductor), and each sample was normalized to the amount of total protein. Raw data are available for download at DOI 10.5281/zenodo.7932681.

### Data analysis and statistics

Unless otherwise noted, data analysis was performed in MATLAB (version R2019b) using built-in functions. Code for generating dose response curves, death kinetic curves, and GRADE plots in MATLAB is available on GitHub (https://github.com/MJLee-Lab). Violin and beeswarm-style plots were generated in GraphPad Prism 9 (version 9.5.0). Pair-wise statistical comparisons were made using a two-sample t-test unless otherwise noted. For the L2FC, growth rate, and death rate values from the chemo-genetic screens, significance was determined empirically by bootstrapping a p-value and performing FDR correction with the Benjamini-Hochberg procedure. GSEA analyses were performed using the GSEA 4.1.0 package, and the associated data was plotted in MATLAB. ImageStudio 4.0.21 was used to analyze western blots. Flow cytometry analysis was performed using FlowJo version 10.8.1. Image analysis was performed using Fiji (ImageJ2, version 2.3.0).

## ACKNOWLEDGEMENTS

We thank current and past members of the UMass Chan Medical School DSB community for their helpful comments and critiques during the design and execution of this study. Additionally, we thank Thomas Leete for his assistance with training in an early stage of this project; Christina Baer and UMass Chan SCOPE Core for assistance with some microscopy experiments; Amir Mitchell for providing an H2B-mCherry plasmid; Thomas Fazzio for providing the pX330 plasmid; Tina Fortier, Eric Baehrecke and the UMass Chan Electron Microscopy Core for assistance with TEM experiments; and Michael Green and Dohoon Kim for providing access to some of the cell lines used in this study. MSI is supported by the Novo Nordisk Foundation Center for Basic Metabolic Research, an independent research center based at the University of Copenhagen, and partially funded by an unconditional donation from the Novo Nordisk Foundation (grant number NNF18CC0034900). This work was supported by grants from the National Institutes of Health/NIGMS (R01 GM127559 to MJL), the NCI (F31 CA268847 to MEH), and the American Cancer Society (RSG-17-011-01 to MJL).

## AUTHOR CONTRIBUTIONS

This project was conceived by MEH and MJL. Chemo-genetic screens, siRNA screens, flow cytometry, and cobalt-calcein assays were performed and analyzed by MEH. Drug sensitivity screens were performed by MEH with assistance from PCG. Cell counting experiments were performed by MEH and REF. GRADE-based evaluation of growth and death rates was performed by MEH and MJL. Imaging experiments were performed and analyzed by MEH, with assistance from SAP. Evaluation of error between chemo-genetic screen analysis strategies was performed by NWH. MSI, DAG, and JBS consulted on the design, interpretation, and analysis of assays to interpret mitochondrial function. BH3 profiling experiments were designed, executed, and analyzed by CSF and KAS. Metabolomic profiling experiments were designed, performed, and analyzed by MEH, in consultation with JBS and MJL. Meta-analysis of published screens was performed by MJL. All other experiments, statistical analyses, and modeling were conducted by MEH. MEH and MJL wrote and edited the manuscript.

## SUPPLEMENTAL FIGURE LEGENDS

**Supplemental Figure 1:**
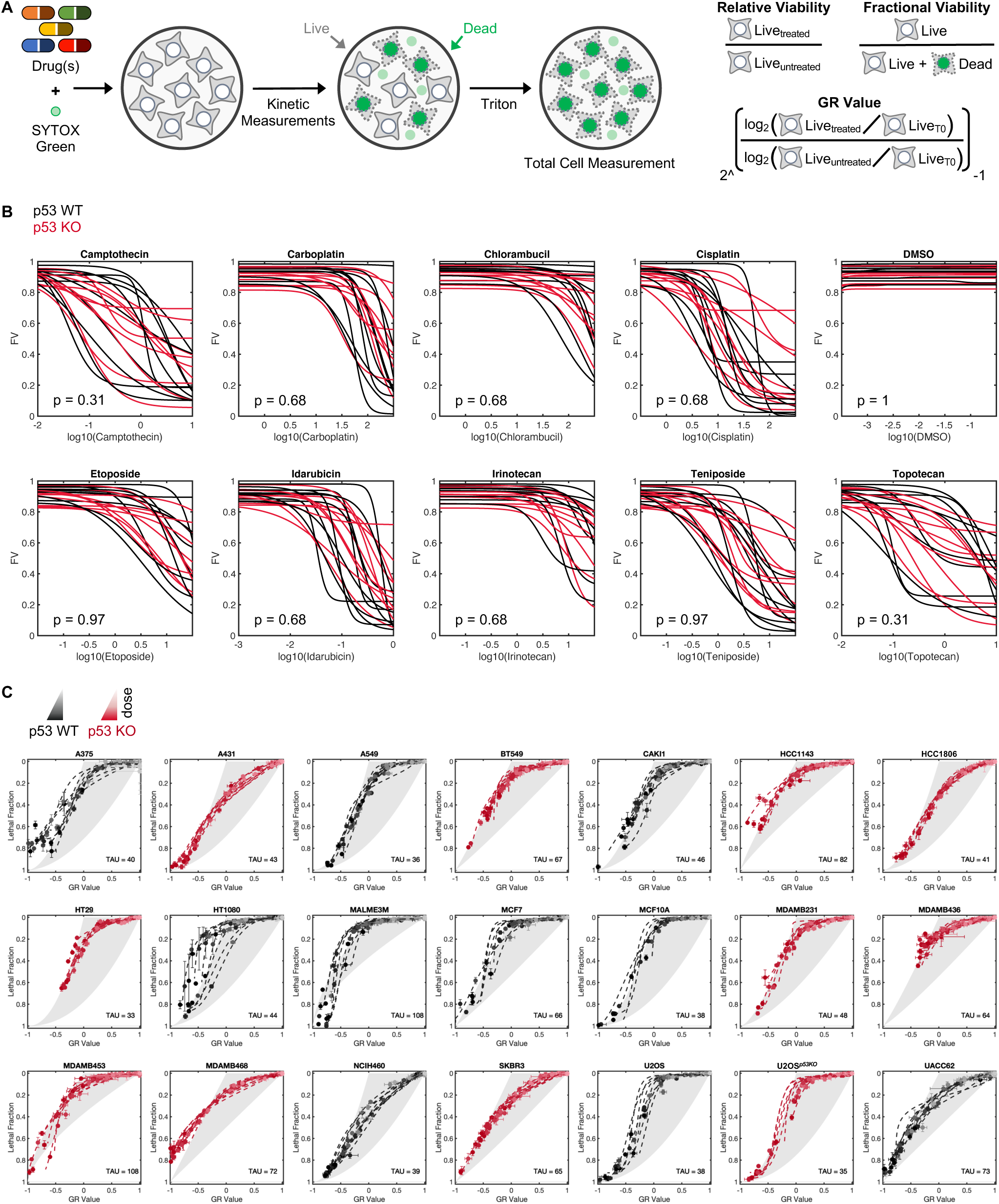
Fractional viability and drug GRADE across p53-proficient and p53-deficient cell lines. Related to Figure 1. (**A**) Schematic of the FLICK assay and equations for calculating relative viability (RV), fractional viability (FV), and GR values. (**B – C**) Sensitivity of p53 WT and p53 KO cell lines to DNA-damaging chemotherapeutics, as measured by (B) FV or (C) drug GRADE.

**Supplemental Figure 2:**
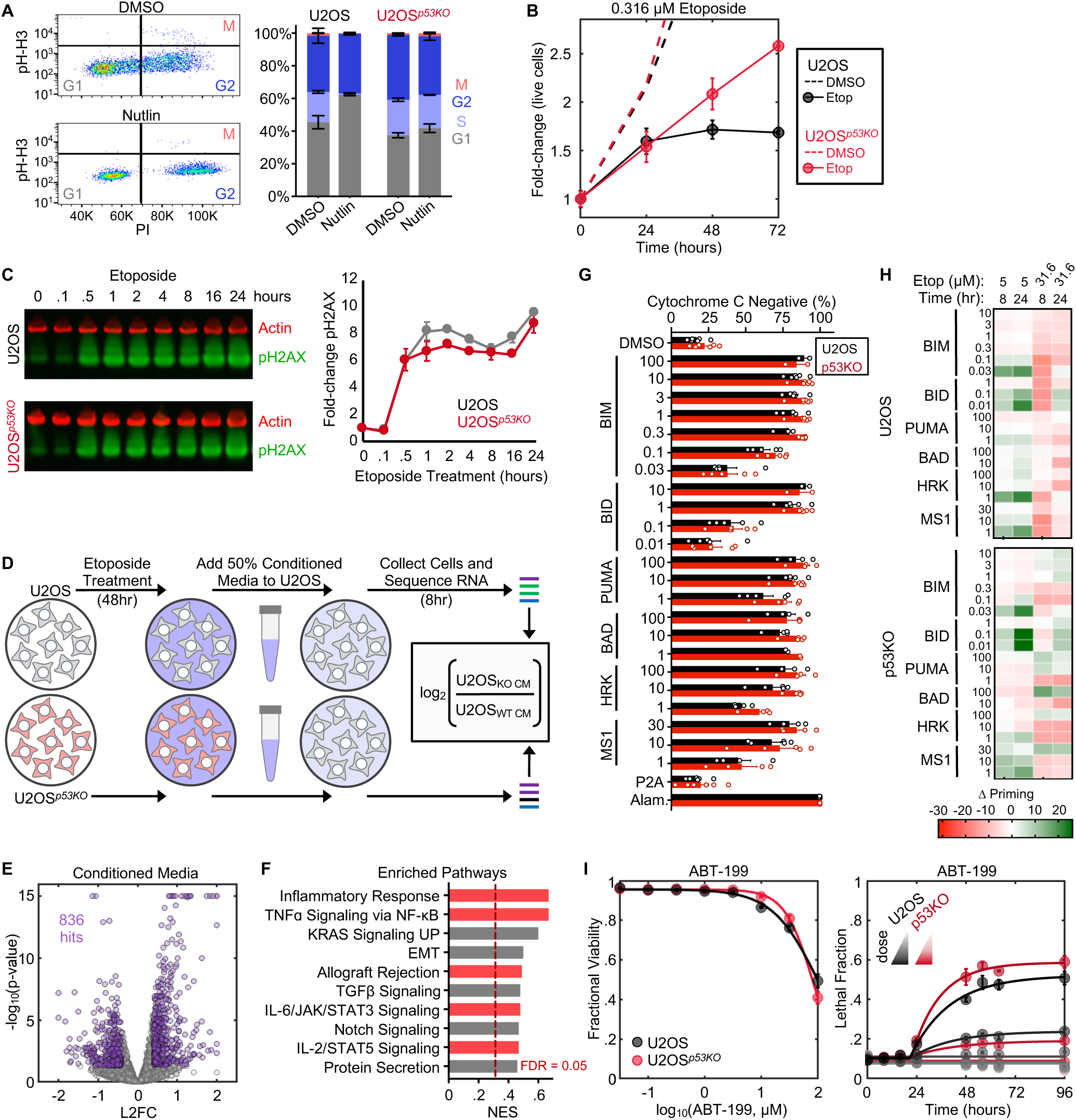
p53 deletion compromises cell cycle arrest but does not prevent activation of DNA repair or BH3 mimetic-induced apoptosis. Related to Figure 1 **and** Figure 2. **(A)** Measurement of cell cycle position using PI staining and the mitotic marker pH-H3. Example for untreated U2OS cells (*left*), and quantification of cell cycle phase from cells treated with nutlin (*right*). **(B)** Live cell counts over time for U2OS and U2OS*^p53KO^* cells treated with a sub-lethal dose of etoposide. **(C)** Kinetic western showing the phosphorylation of the DNA damage marker H2AX (phospho-S139) in response to etoposide. Quantification ± SD from 3 experimental replicates (*right*). **(D-F)** Evaluation of inflammatory cell death in U2OS and U2OS*^p53KO^*cells. **(D)** Schematic for conditioned media experiment. **(E)** Volcano plot showing the p-values and L2FCs for U2OS cells treated with conditioned media (log_2_(U2OS*^p53KO^*/U2OS)). **(F)** Pathway-level enrichment for conditioned media, highlighting enrichment for inflammatory signatures in cells treated with media conditioned by U2OS*^p53KO^* cells. **(G-H)** Apoptotic priming evaluated using BH3 profiling. **(G)** Basal BH3 profiles in U2OS and U2OS*^p53KO^* cells. **(H)** Change in apoptotic priming level following etoposide exposure. **(I)** Activation of apoptotic death in U2OS and U2OS*^p53KO^*cells by the BH3 mimetic ABT-199.

**Supplemental Figure 3:**
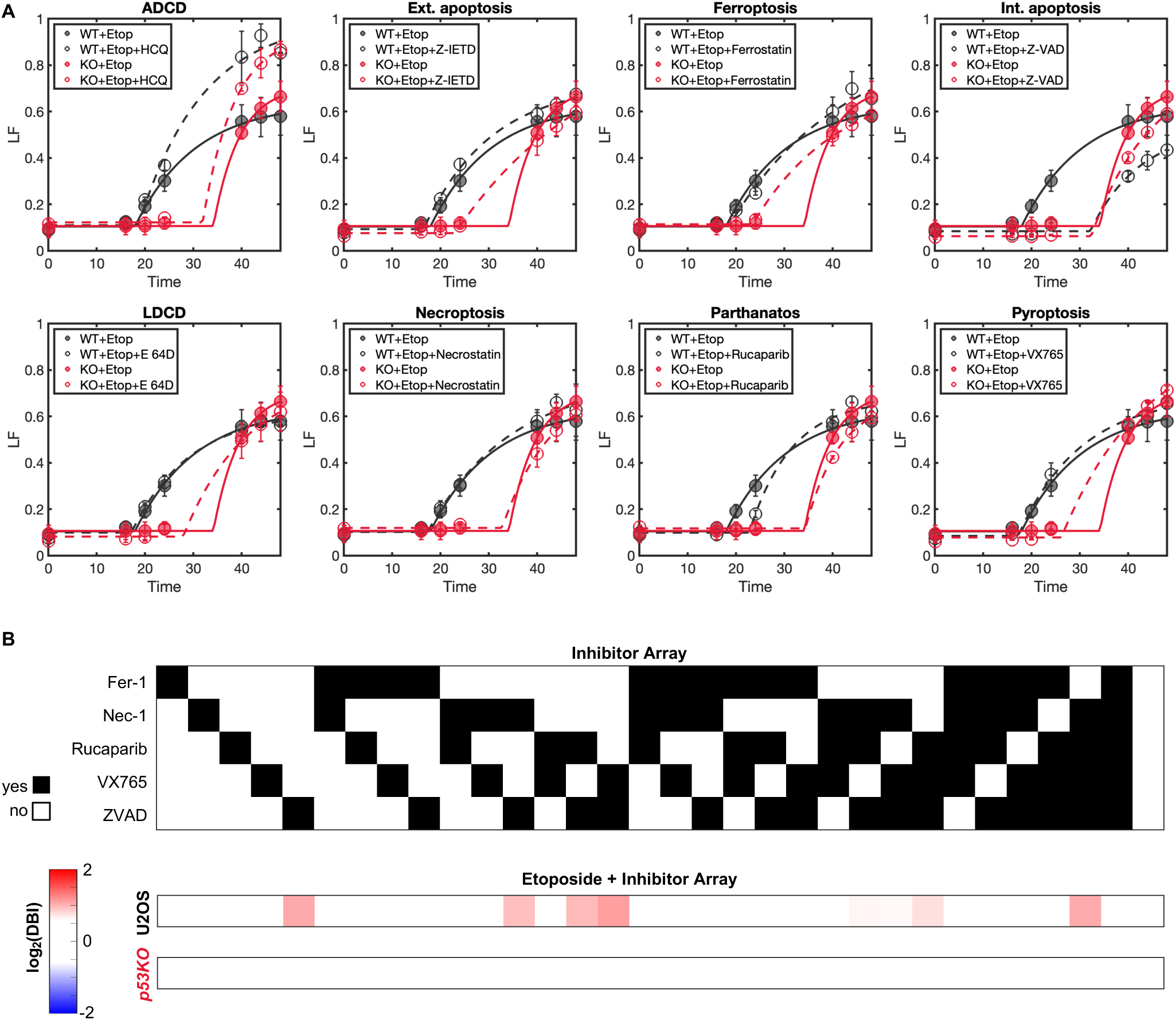
U2OS and U2OS*^p53KO^* treated with cell death inhibitors. Related to Figure 3. **(A)** U2OS and U2OS*^p53KO^* cells treated with single inhibitors for 8 common cell death pathways. **(B)** U2OS and U2OS*^p53KO^* cells treated with higher-order combinations of 5 cell death inhibitors. Heatmap colored by Deviation from Bliss Independence (DBI). Negative DBI values report enhanced lethality, positive DBI reports inhibition of lethality.

**Supplemental Figure 4:**
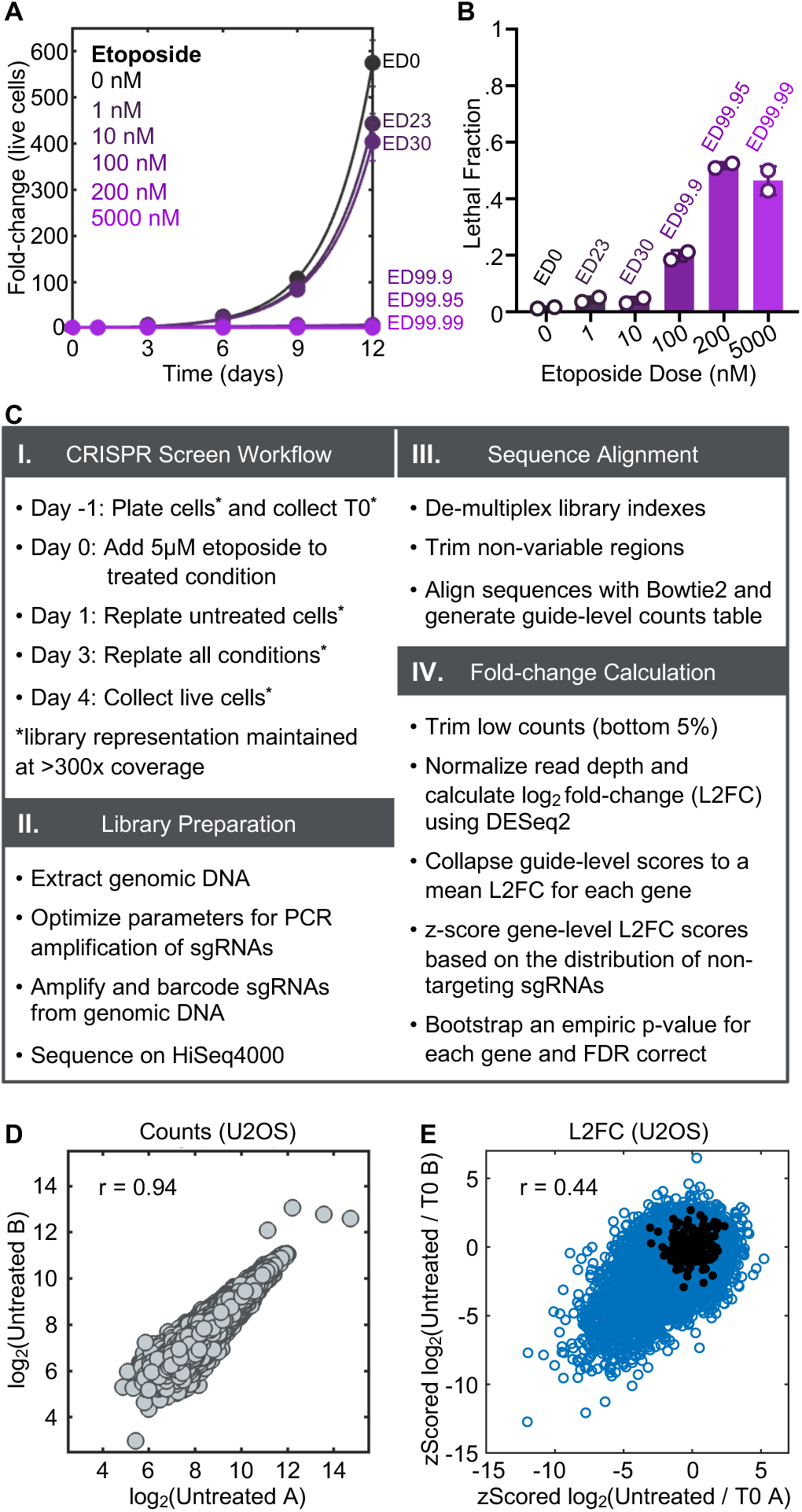
Chemo-genetic screening analysis strategy and replicate correlation. Related to Figure 3. **(A – B)** U2OS cells treated with etoposide for 12 days. (A) Live cells were counted to determine the growth defect of each dose. ED = “Effective Dose” (e.g., ED30 = effective dose for 30% reduction in population size after 12 days, compared to untreated). (B) Dead and live cells were counted to determine fractional viability at each dose. **(C)** Analysis schematic for calculating L2FC from chemo-genetic screens. **(D)** Example of correlation between counts for two replicates of the same screen condition. **(E)** Example of correlation between gene-level L2FC values for two screen replicates. z scored L2FC was calculated for each replicate.

**Supplemental Figure 5:**
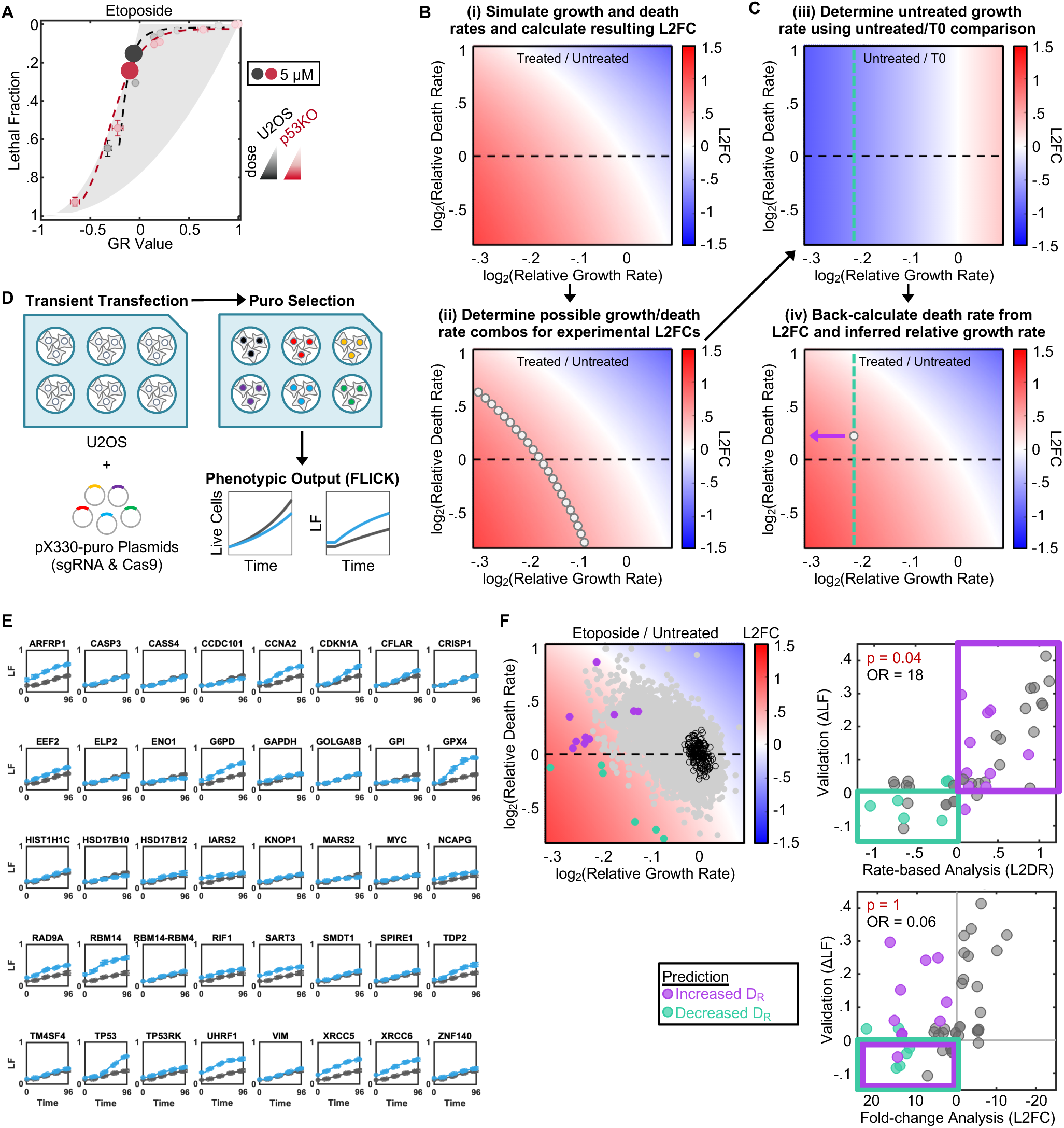
Validation of rate-based analysis method for chemo-genetic screen. Related to Figure 4. **(A)** Drug GRADE for U2OS and U2OS*^p53KO^* cells treated with etoposide. Dose selected for the CRISPR screen (5 μM) is highlighted. **(B – C)** Schematic for calculation of drug-induced death rate and growth rate from experimentally observed L2FC values. (B) Phase diagram and scatter to highlight one example L2FC that can be produced from multiple combinations of growth and death rate. (C) Calculation of growth rate and inference of the drug-induced death rate. **(D)** Schematic of method used to validate hits from the whole-genome CRISPR screen. **(E)** Validation data generated using FLICK, black = non-targeting sgRNA, blue = targeted gene. **(F)** Phase diagram and scatter plots highlighting validated genes that have a reduced growth rate and are predicted to induce resistance using L2FC. p-values and odds ratios (OR) calculated using a Fishers exact test.

**Supplemental Figure 6:**
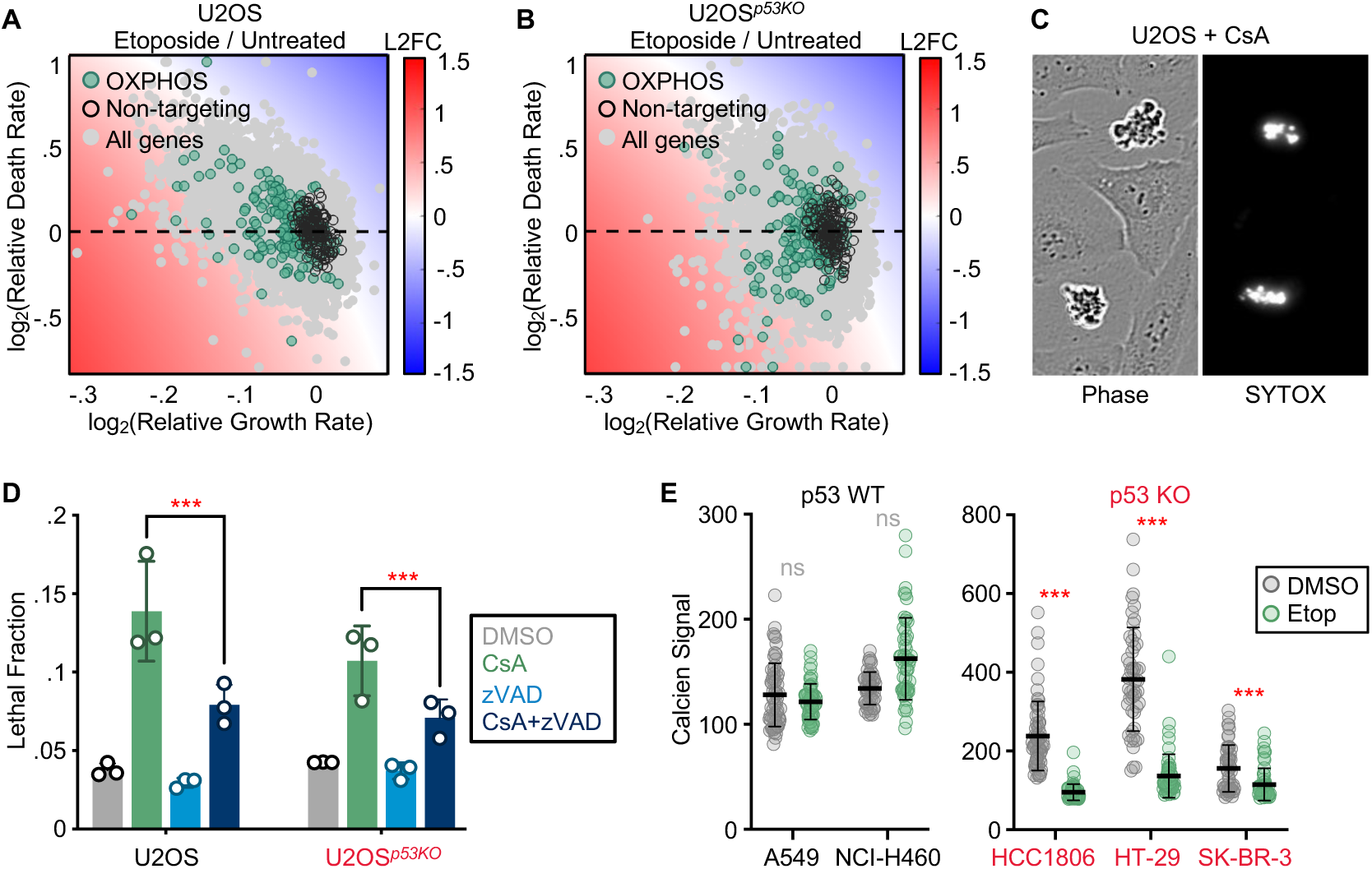
Validation of MPT activation in p53 KO cells. Related to Figure 5. **(A)** Gene-level chemo-genetic profiling data for U2OS cells, projected into phase diagram, with OXPHOS genes highlighted. **(B)** as in (A), but for U2OS*^p53KO^* cells. **(C)** Phase and SYTOX green images of U2OS cells treated with 10 μM cyclosporin A (CsA). **(D)** Lethal fraction of U2OS and U2OS*^p53KO^*cells treated with CsA, zVAD, or CsA+zVAD. **(E)** Cobalt-calcein assay performed on naturally p53-proficient and p53-deficient cell lines treated with 31.6 μM etoposide for 36 hours.

**Supplemental Figure 7:**
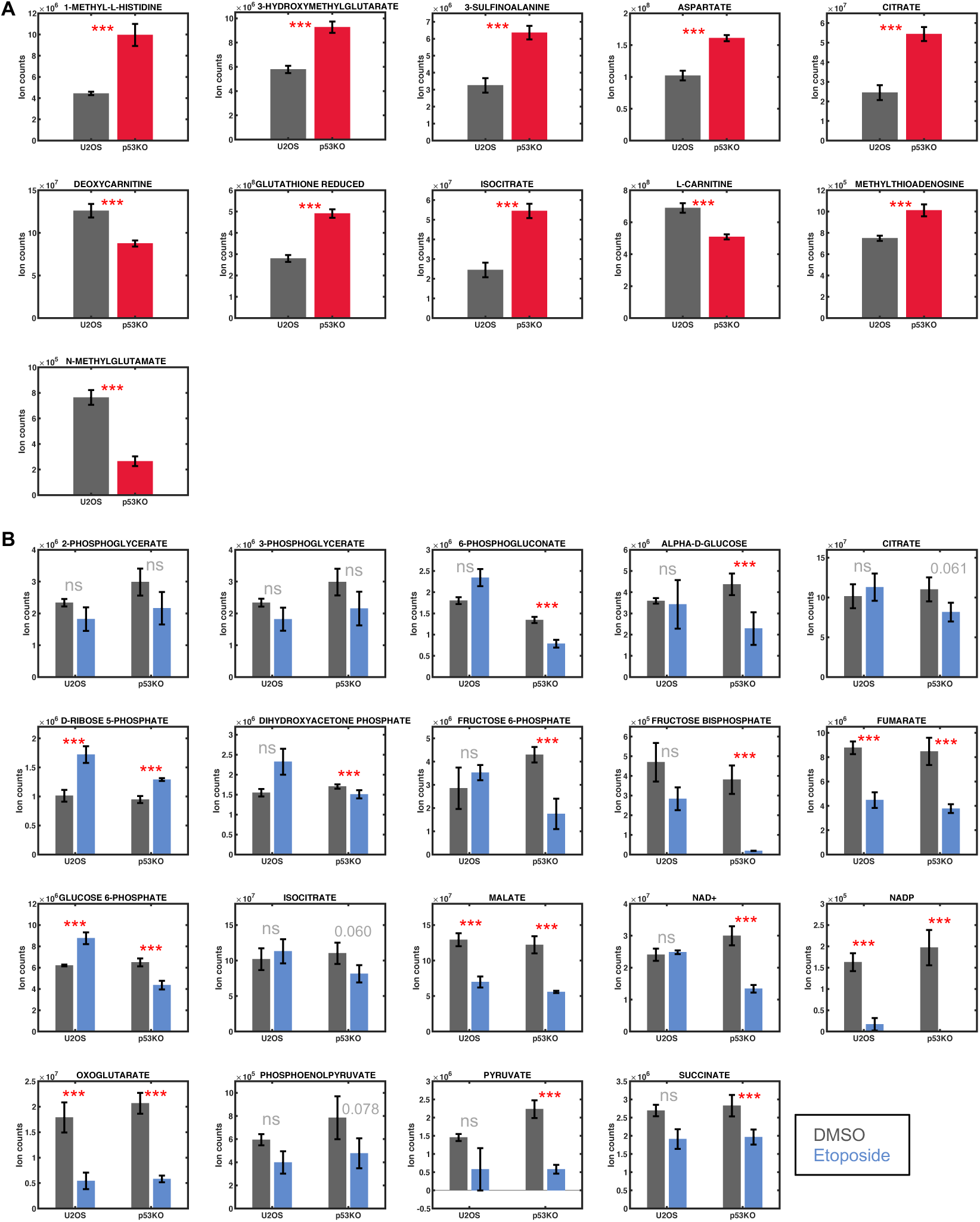
Steady state metabolite levels. Related to Figure 7. **(A)** Metabolite levels shown for vehicle treated U2OS and U2OS*^p53KO^*cells at T0. **(B)** Metabolite levels shown for intermediate metabolites involved in glycolysis, PPP, or TCA cycle for U2OS and U2OS*^p53KO^*. Data are shown for vehicle and etoposide treated samples at 48 hours. Asterisks report p<0.05. ns = not significant. P-values highlighted for metabolites that were not significant at 48 hours but were significantly different at 24 hours. Data are mean ± SD from 3 experimental replicates.

**Supplemental Figure 8:**
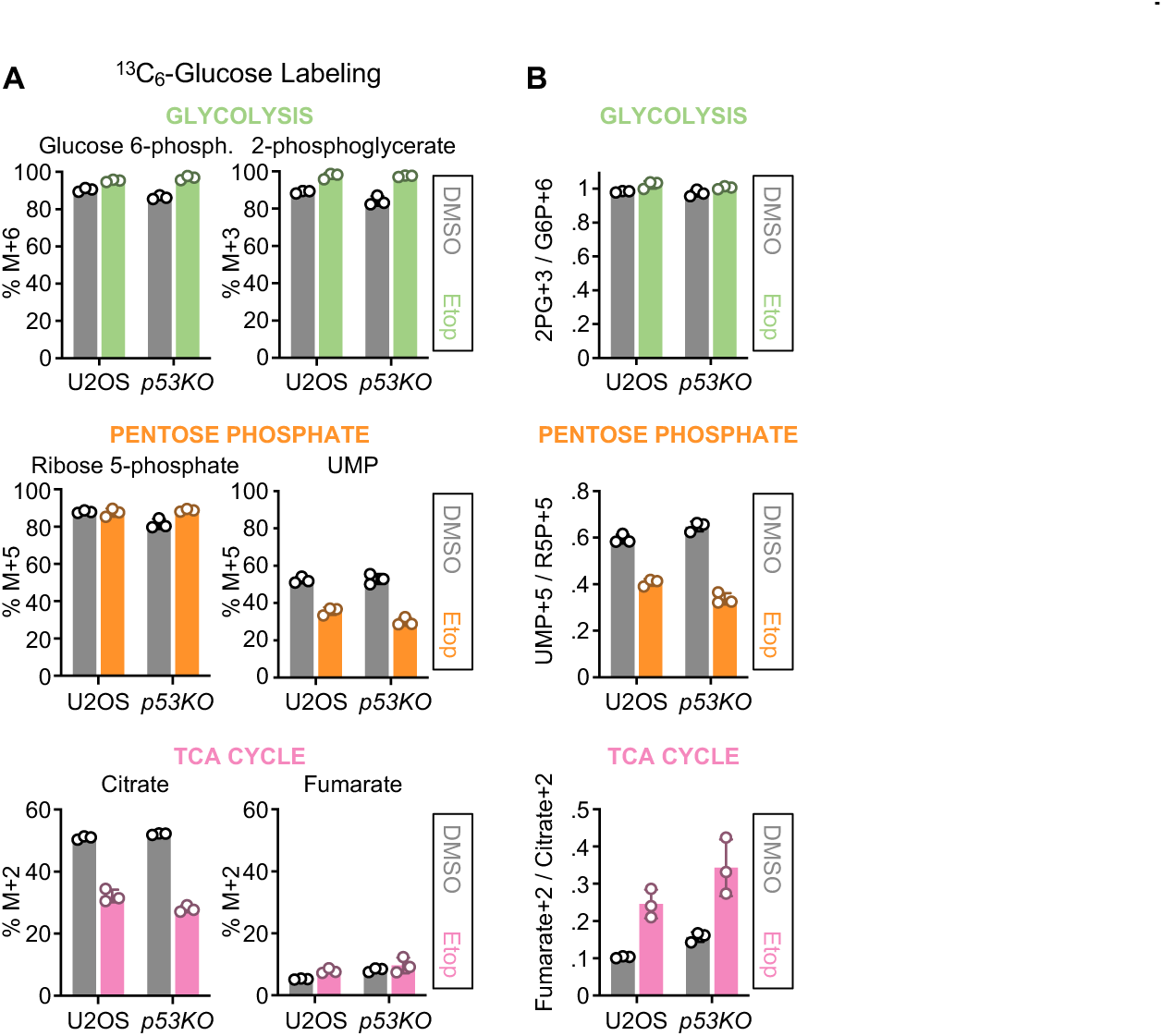
Proportional enrichment of glucose-derived metabolites is not altered by the loss of p53. **(A)** Isotope tracing using ^13^C_6_-Glucose. Cells were labeled for 8 hours. Fractional enrichment for intermediate metabolites in glycolysis, PPP, and TCA cycle shown for U2OS and U2OS*^p53KO^* cells treated with etoposide for 48 hours. **(B)** Data collected as in (A) but analyzed to compare the fractional enrichment of glucose-derived metabolites for upstream and downstream components of glycolysis, PPP, or the TCA cycle. Abbreviations: glucose 6-phosphate (G6P), 2-phosphoglycerate (2PG), ribose 5-phosphate (R5P), uridine monophosphate (UMP).

